# Sacituzumab Govitecan as an Effective Strategy for Sensitizing Chemoresistant HNSCC Cells to Senolytic Intervention

**DOI:** 10.64898/2026.04.13.718209

**Authors:** Natalie Luffman, Bin Hu, Jennifer Koblinski, David A. Gewirtz, Hisashi Harada

**Affiliations:** Department of Cellular, Molecular, and Genetic Medicine, School of Medicine; Philips Institute for Oral Health Research, School of Dentistry; Massey Comprehensive Cancer Center; Virginia Commonwealth University, Richmond, VA; Cancer Mouse Models Core, Massey Comprehensive Cancer Center; Virginia Commonwealth University, Richmond, VA; Department of Pathology, School of Medicine; Cancer Mouse Models Core, Massey Comprehensive Cancer Center; Virginia Commonwealth University, Richmond, VA; Department of Pharmacology and Toxicology, School of Medicine; Massey Comprehensive Cancer Center; Virginia Commonwealth University, Richmond, VA; Philips Institute for Oral Health Research, School of Dentistry; Massey Comprehensive Cancer Center; Virginia Commonwealth University, Richmond, VA

## Abstract

Head and neck squamous cell carcinoma (HNSCC) is the sixth most prevalent cancer worldwide and is marked by a high tumor relapse due to acquired chemoresistance, requiring alternative strategies to sensitize resistant tumors to treatment. Sacituzumab govitecan (SG), a TROP2-targeting antibody-drug conjugate, has been successful in limiting tumor progression in pretreated patients with triple-negative and hormone-receptor positive HER2-negative breast cancer. However, it has been ineffective as a monotherapy in HNSCC. This may be attributed to the promotion of senescence ultimately leading to tumor relapse. We established that isogenic cisplatin-sensitive and -resistant HNSCC cells undergo senescence following SG treatment. Senolytics have been effective in sensitizing solid tumors to standard of care chemotherapies in preclinical studies. Consequently, we investigated the effectiveness of SG treatment followed by the senolytic, ABT-263, as a “two-hit” therapeutic strategy against cisplatin-resistant HNSCC. SG followed by ABT-263 sensitized HNSCC cells to apoptosis via a BAK and BAX-dependent mechanism. *In vivo* studies confirmed that SG treatment followed by ABT-263 limited tumor progression and extended survival without notable toxicity. Thus, SG in combination with senolytic treatment may be an effective strategy for overcoming not only cisplatin-resistance but also SG resistance via targeting the senescent cell population.

## Introduction

DNA-damaging platinum-based chemotherapeutic agents such as cisplatin are the current standard of care therapies for head and neck cancer (HNC) patients and are typically applied in combination with radiation^1,2^. High doses of cisplatin are administered over the course of several weeks, though toxicities may reduce the frequency of doses^2,3^. Alternatively, multiple rounds of weekly low-dose cisplatin infusions may be administered over several months in conjunction with radiation^3^. Cisplatin binds to nuclear DNA to create inter- and intra-strand crosslinks between purine residues, inducing single-stranded DNA breaks that ultimately trigger downstream cell death pathways^4^. Though the majority of HNC patients are initially responsive to cisplatin, approximately 50% of HNC patients will experience tumor relapse, caused by chemoresistant tumors that are no longer responsive to cisplatin treatment^1,2^. This may be a result of upregulation of DNA damage repair pathways, alterations in drug uptake and export following continuous cisplatin exposure^1^, and the promotion of senescence^5,6^. As a result, these resistant tumors are no longer responsive to conventional therapeutics and require alternative treatment strategies to overcome chemoresistance while mitigating toxicities.

Antibody-drug conjugates (ADCs) are an emerging drug class being tested in a variety of solid tumors^7,8^. ADCs are comprised of a human monoclonal antibody targeting a tumor-specific antigen, a chemotherapeutic warhead, and a linker between the warhead and the antibody^7,9^. The tumor-specific antibody allows efficient and targeted delivery of the cytotoxic payload to the tumor cell population, thus limiting off-target effects in healthy, non-cancerous cells^7,9^. Once bound to the designated antigen, the ADC is endocytosed by the cell and the linker is cleaved, releasing the payload into the cytosol and promoting cell death^7,9^.

Sacituzumab govitecan (SG), is a third-generation TROP2-targeting ADC delivering the topoisomerase I inhibitor SN-38 (irinotecan). TROP2 is a transmembrane protein regulating calcium signaling, cell proliferation, and cell cycle progression and has been found to be overexpressed in a variety of solid tumors, including breast, pancreatic, and head and neck^10–12^. A CL2A linker conjugates the anti-TROP2 antibody (sacituzumab) and SN-38 warhead, allowing SG to induce cell death via DNA damage from inhibition of topoisomerase I^13,14^. It is currently approved by the FDA for metastatic triple negative breast cancer (mTNBC) and hormone-receptor positive HER2-negative breast cancer patients who have received previous rounds of systemic chemotherapies^15,16^. Furthermore, SG has been found to successfully reduce tumor growth in platinum-resistant ovarian and urothelial cancer^17,18^. Datopotamab deruxtecan (Dato-DXd), an additional TROP-2 targeting ADC similarly inhibiting topoisomerase I, has also been clinically successful in limiting tumor progression and reducing tumor burden in pretreated non-small cell lung cancer patients^19,20^. Thus, TROP2-targeting ADCs have proved to be effective adjuvant therapies for solid-tumor patients while potentially overcoming acquired resistance from previously treated standard chemotherapies and limiting overall toxicities.

Despite success in breast cancer, a phase II basket trial indicated that SG has little impact as a single agent in head and neck squamous cell carcinoma (HNSCC) patients previously receiving cisplatin or anti-PD-1 therapy^12,21^. This moderate efficacy in head and neck cancer may partly be due to therapy-induced senescence within the tumor cell population, where cells become dormant following chemotherapy yet continue to be metabolically active. As a result, senescent cells may acquire chemoresistance, eventually recovering their proliferative capacity leading to tumor relapse^22,23^. Senescent cells additionally upregulate an array of secreted proteins, including cytokines, chemokines, and growth factors, collectively known as the senescence associated secretory phenotype (SASP). These secreted factors may regulate autocrine and paracrine signaling within the tumor microenvironment that may stimulate tumorigenesis, immune evasion, and epithelial-mesenchymal transition, leading to acquired chemoresistance and aggressive tumor relapse^23–25^. Prolonged treatment with SG may result in decreased SG sensitivity promoting ADC resistance^26^.

Senolytics, drugs specifically inducing cell death in senescent cell populations, have emerged as an effective strategy for eliminating senescent cells via inhibiting anti-apoptotic BCL-2 family proteins that are often upregulated following senescence induction^23,27^. ABT-263, a BCL-X_L_/BCL-2 inhibitor, has been found to selectively target senescent cells in ovarian, breast, prostate, and head and neck cancer via targeting BCL-X_L_, thus mitigating tumor progression *in vitro* and *in vivo*^28–30^. Here we demonstrate that SG is capable of inducing senescence in cisplatin-sensitive and -resistant HNSCC cells. Furthermore, we show that a two-hit strategy of SG treatment followed by ABT-263 could be effective against SG-induced senescence. By targeting and reducing the senescent cell population, this combination strategy of SG paired with senolytics may further limit acquired resistance and promote antitumor activity.

## Materials & Methods

### Cell culture & drug sensitivity

HPV-negative human HN30, HN30R, Detroit-562, and Detroit-562R cells were maintained in 1x Dulbecco’s Modified Eagle Medium (1x DMEM; Thermo Fischer, Waltham, MA USA) supplemented with 100 U/mL penicillin G sodium (Thermo Fischer), 100 µg/mL streptomycin sulfate, and 10% fetal bovine serum (Thermo Fischer). HN30R cells were further supplemented with 5 μM cisplatin while Detroit-562R cells were supplemented with 2.5 μM cisplatin (MedchemExpress, South Brunswick, NJ, USA, HY-17394). All cells were housed in an incubator at 5% CO_2_ and 37°C. Cisplatin was dissolved in 1x PBS while Sacituzumab Govitecan (SG; Trodelvy, NDC 5513513201) and a naked sacituzumab antibody (MedchemExpress, HY-P99045) were suspended in 0.9% saline. SN-38 (MedchemExpress, HY-13704), ABT-263 (MedchemExpress, HY-10087), S63845 (MedchemExpress, HY-100741), A-1155463 (MedchemExpress, HY-19725), and ABT-199 (MedchemExpress, HY-15531) were dissolved in DMSO. HN30 cells were originally provided by Dr. Andrew Yeudall (Augusta University, Augusta, GA, USA).

Cisplatin, SG, and SN-38 sensitivities were evaluated via WST-1 assay to identify the IC_50_ concentrations. Cells were seeded in 96-well plates and treated with either 0-100 μM cisplatin, 0-10 μg/mL SG, or 0-0.05 μM SN-38 for 72 hours followed by treatment with WST-1 (MilliporeSigma, St. Louis, MO, USA, 5015944001) for 2-3 hours at 5% CO_2_ and 37°C. Plates were then assessed for spectrophotometric absorbance at 450 nm via the Glomax multi-detection system (Promega, Madison, WI, USA).

### qRT-PCR

Cells were treated with the desired concentration of SG for 72 hours and harvested for RNA isolation via ZymoPure RNA mini-prep kit (Zymo Research, Irvine, CA, USA, R1054). cDNA was then prepared using the Applied Biosystems cDNA kit (Applied Biosystems, Foster City, CA, USA, 4368814). Prepared cDNA was then used for qPCR analysis using the Applied Biosystems SYBR Green reagent (Applied Biosystems, A46109) and the human primers in **Supplementary Table 1** synthesized via Integrated DNA Technologies (Coralville, IA, USA).

### Senescence-associated β-galactosidase (SA-β-gal) analyses

*In vitro* histochemical staining of β-galactosidase was performed using the X-gal detection kit (Cell Signaling, Danvers, MA, USA, 9860). Cells were seeded in 6-well plates and fixed with 1x fixation buffer for 10-15 minutes followed by X-gal staining for 24 hours. Cells were then imaged via bright field inverted microscope (Olympus, Tokyo, Japan).

*Ex vivo* β-galactosidase staining was implemented in HN30R tumors harvested from NSG mice following treatment with either vehicle, SG-only, ABT-263-only, or SG followed by ABT-263. Vehicle and SG-only treated tumors were harvested ten days after SG treatment, ABT-263-only treated tumors were collected three days after ABT-263 treatment, and SG+ABT-263 treated tumors were harvested five days after ABT-263 treatment. Dissected tumors were embedded in OCT cryopreservation medium (Sakura Finetek, Torrance, CA, USA, 4583) on dry ice and submitted for 10 μm sectioning. Slide-mounted tumors were then fixed in a solution of 0.2% formaldehyde and 0.02% glutaraldehyde in 1x PBS for 10 minutes and subsequently washed in 1x PBS before staining. Slides were then stained in an X-gal solution (40 mM citric acid/sodium-phosphate buffer, 5 mM potassium ferricyanide, 150 mM sodium chloride, and 2 mM magnesium chloride in DI H_2_O) overnight at 37°C in a CO_2_-free incubator. Slides were then washed in DI H_2_O and then dehydrated via five-minute subsequent washes in increasing concentrations of 50-100% 200-proof ethanol followed by a final wash in isopropanol for 5 minutes. Permount (Fischer Scientific, Waltham, MA, USA, SP15-100) was then added to cover each tumor section before coverslip mounting. Tumor slides were imaged via bright field inverted microscope (Olympus, Tokyo, Japan) at 10x magnification.

DDAOG β-gal FACS analysis was utilized to observe senescence induction following SG or SN-38 treatment ^31^. Cells were seeded and treated as described and then harvested after 72 hours. Cells were incubated in a 1:1000 1x DMEM-Bafilomycin (MedchemExpress, HY-100558) solution at 37°C for 30 minutes followed by DDAOG (Invitrogen, Carlsbad, CA, USA, D6488) staining at a 1:1000 dilution for 1 hour. Samples were washed three times in ice-cold 0.5% BSA-PBS and then resuspended in 1% BSA-PBS before sorting via the BD LSRFortessa-X20 cytometry at the Virginia Commonwealth University (VCU) Flow Cytometry Core.

### Western blotting

Cells were treated at the desired concentrations with drugs and then harvested for total cell lysates at the indicated timepoints. Western blotting was performed as described previously^30^. The following primary antibodies were implemented at a 1:1000 dilution: GAPDH (Proteintech, Rosemont, IL, USA, 60004-1-Ig), p21 (Cell Signaling, 2947), BCL-X_L_ (Cell Signaling, 2764), BAK (Cell Signaling, 14155), BAX (Cell Signaling, 5023), Cleaved-Caspase 3 (Cell Signaling, 9664), and TROP2 (Cell Signaling, 47866). The following secondary antibodies were implemented at a 1:2000 dilution: anti-rabbit IgG HRP-linked (Cell Signaling, 7074), anti-mouse IgG HRP-linked (Cell Signaling, 7076). Membranes were imaged on Chemidoc (Bio-Rad, 12003153) with automatic exposure. ImageJ 1.53k was utilized to calculate the densitometry for each blot indicated. N/Tert-1 keratinocyte lysates were provided by Dr. Molly Bristol (VCU)^32^.

### Generation of knockdown cell lines with shRNA

Plasmid shRNA constructs shTROP2 #1 (TRCN0000056419), shTROP2 #2 (TRCN0000056421), shBAK (TRCN0000033466), shBAX (TRCN0000312625, used for shBAX single knockdown, puromycin-resistant were purchased from Sigma Aldrich (St. Louis, MO); shBAX (HSH150291-LVRU6H-c, used for shBAK/shBAX double knockdown, hygromycin-resistant) were purchased from GeneCopoeia (Rockville, MD, USA) while a scrambled RNA control was purchased from Addgene (Watertown, MA, USA, 1864). Knockdown plasmids were transfected into HEK293T cells (ATCC, Manassas, VA; RRID: CVCL_0063) using Endofectin (GeneCopeia, EF001) with psPAX2 (RRID: Addgene_12260) and pMD2.G (RRID: Addgene_12259) plasmids to generate their respective lentiviruses. HN30 and HN30R cells were infected with the desired lentiviruses and continuously treated with 2 μg/mL puromycin (Invivogen, San Diego, CA, USA, ant-pr) or 0.1 μg/mL hygromycin (GoldBio, St. Louis, MO, USA, H-270-1) to select for shRNA expressing cells.

### Cell death analysis

Cell viability was qualitatively observed via Crystal-Violet staining. Cells were seeded in 6-well plates and treated with either 0.9% saline vehicle or SG followed by treatment with either DMSO, ABT-263, S63845, A-1155463, or ABT-199. Cells were then washed with 1x PBS followed by fixation with methanol for 15 minutes. Fixed cells were stained with 0.05% crystal-violet solution (Thermo Fischer, 525275B) and washed with DI water three times.

Apoptosis was quantitatively assessed via Annexin-V/PI flow cytometry. Cells were seeded and treated with the indicated concentrations of SG and ABT-263 and harvested at the desired timepoint. Collected cells were washed in 1x PBS, resuspended in 100 µL 1x binding buffer, and then stained with Annexin-FITC (Biolegend, San Diego, CA, USA, 640945) and propidium iodide (Invitrogen, P3566) and incubated in the dark for 15 minutes. Then 400 µL 1x binding buffer were added to the final sample suspensions. The BD LSRFortessa-X20 cytometer was used to analyze the samples via the VCU Flow Cytometry Core.

### Proliferative Recovery Assay

Trypan blue exclusion assay was assessed for live cell count at the indicated timepoints during and after SG ± ABT-263 treatment. Cells were collected via 0.25% trypsin-EDTA treatment followed by staining with 0.4% trypan blue (Thermo Fischer, 15250-061). Collected cells were then counted via TC20 Automated Cell Counter (BioRad, Hercules, CA, USA, 1450102).

### In vivo studies

The animal study was performed according to the VCU Institutional Animal Care and Use Committee guidelines (VCU IACUC Protocol #AD1001436). 6-8 week-old male (HN30R cells were originally established from a male patient) NOD-scid-IL2Rγ-null (NSG) mice (Fig. 6 study: *n = 24*; Supplemental Fig. 3 study: *n* = 32) were provided by the VCU Cancer Mouse Models Core. HN30R cells were suspended in a 1:1 PBS to basement membrane extract (BME, Matrigel type 3) solution (Biotechne, Cat. # 3631-010-02) for tumor inoculation. HN30R cells (1x10^6^) were injected into the left cheek of each mouse (Day 0)^33^. Mice were monitored 3x weekly until tumors reached approximately 50 mm^3^ in volume where all mice were randomized into four treatment groups (*n* = 6-8 per group): vehicle (0.9% saline), SG (0.25 mg/kg), ABT-263 (80 mg/kg), and SG+ABT-263. SG was administered via intraperitoneal injection while ABT-263 was given via oral gavage at the indicated timepoints. Tumor volume was assessed via caliper measurements where volume was calculated by *V =* 0.5*(*l*\*(*w^2^*)), (*l =* tumor length, *w* = width, and *l* > *w*). Mice were regularly given 76A DietGel (ClearH2O; Portland, ME; 72-07-5022X) following SG and ABT-263 treatment to limit weight loss. Mice were humanely euthanized just before tumors reached their humane endpoint of 500 mm^3^ total volume or were significantly ulcerated.

### Statistical analysis

All quantitative data was calculated at the mean ± SEM from a minimum of three independent biological replicates. Microsoft Excel software was used for statistical analysis using an unpaired Student’s t-test.

## Results

### Isogenic cisplatin-sensitive and -resistant HNSCC cells are sensitive to sacituzumab govitecan

To evaluate the efficacy of SG in sensitizing HNSCC cells, we implemented a paired isogenic model of parental cisplatin-sensitive HN30 (male, p53^WT^) and Detroit-562 (female, p53^R175H^) cells and -resistant HN30R and Detroit-562R cells. The HN30R and Detroit-562R cells were generated by continuous treatment of HN30 and Detroit-562 cells with 5 μM and 2.5 μM cisplatin, respectively^34^. The WST-1 assay presented in **Fig. 1A** evaluates the cisplatin sensitivity of both HN30/HN30R and Detroit-562/Detroit-562R cells following 72 hours of treatment. HN30R cells presented with an IC_50_ approximately 3-fold higher (15.5 μM) than HN30 cells (5.0 μM) while Detroit-562R cells displayed an IC_50_ over 2.5-fold higher (20 μM) than Detroit-562 cells (8 μM). **Fig. 1B** indicates that HN30/HN30R and Detroit-562/Detroit-562R cells were both sensitive to SG treatment, with an IC_50_ 0.3 μg/mL and 0.2 μg/mL in HN30 and HN30R cells, respectively, while both Detroit-562 and Detroit-562R cells had an IC_50_ of approximately 1 μg/mL. **Fig. 1C** confirmed that both HN30 and HN30R cells express high levels of TROP2 compared to a keratinocyte control while Detroit-562 and Detroit-562R cells express comparatively lower TROP2 levels. These findings suggest that both cisplatin-sensitive and - resistant HN30 cells are more sensitive to SG treatment potentially due to their high(er) expression of TROP2.

**Figure 1:**
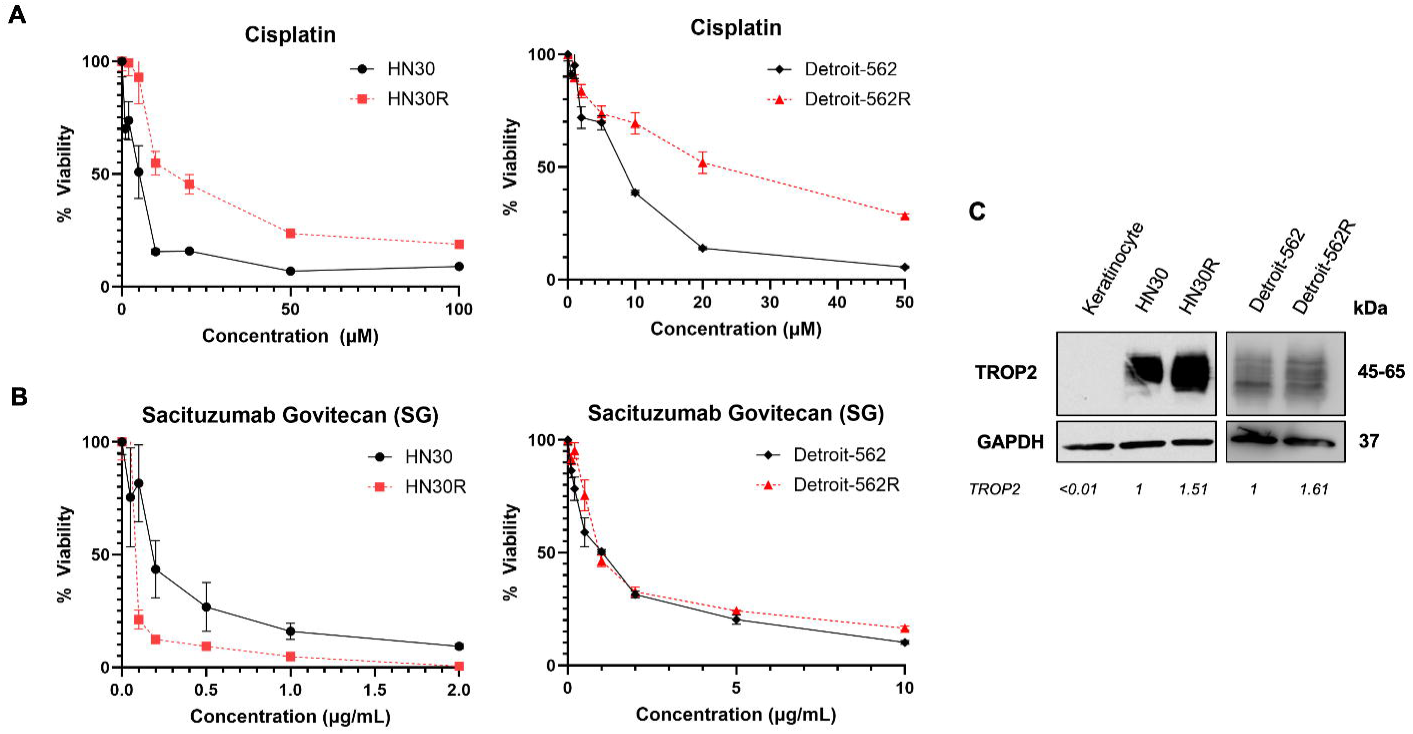
Cisplatin-sensitive and -resistant HNSCC cells are sensitive to SG treatment. **(A)** Cisplatin-sensitive HN30 and Detroit-562 cells and cisplatin-resistant HN30R and Detroit-562R cells were established and treated with cisplatin for 72 hours. Cell viability was assessed via WST-1 assay. Each value represents the average of three biological replicates. **(B)** HN30/HN30R and Detroit-562/Detroit-562R cells were treated with sacituzumab govitecan (SG) for 72 hours and cell viability was determined via WST-1 assay. **(C)** Twenty micrograms of total cell lysates from untreated HN30/HN30R and Detroit-562/Detroit-562R cells were subjected to western blot analysis via the indicated antibodies. Human keratinocytes immortalized by N/TERT were implemented as a negative control. Densitometric quantification was assessed via ImageJ 1.53k; all samples were normalized to GAPDH while fold change was calculated by comparing each sample to the TROP2 level of HN30 cells. Densitometry analysis displays the average of three representative biological replicates.

### SG induces senescence in cisplatin-sensitive and -resistant HNSCC cells

Previous studies from our research groups have reported that SG can induce senescence across multiple solid tumors, including breast, prostate, lung, ovarian, and colorectal cancers^35^. Thus, we evaluated the ability of SG to induce senescence via β-galactosidase activity in both cisplatin-sensitive and -resistant HNSCC cells. HN30 and HN30R cells treated with 0.2 μg/mL SG for 72 hours displayed increased senescence-associated β-galactosidase activity as indicated by histochemical staining of X-gal (**Fig. 2A, left panel**)^36^. FACS analysis assessing DDAOG, a far-red fluorophore released when β-galactosidase becomes active^31^, indicated a significant increase in senescence (35% and 60%) for HN30 and HN30R cells, respectively, following SG treatment (**Fig. 2A, right panel**). Detroit-562 and Detroit-562R cells similarly displayed a 40-60% increase in senescence induction with SG treatment (**Supplemental Figure 1**). Notably, HN30R cells displayed a significantly higher level of senescence induction compared to HN30 cells, which might correlate with higher TROP2 expression in the HN30R cells (**Fig. 1C**).

**Figure 2:**
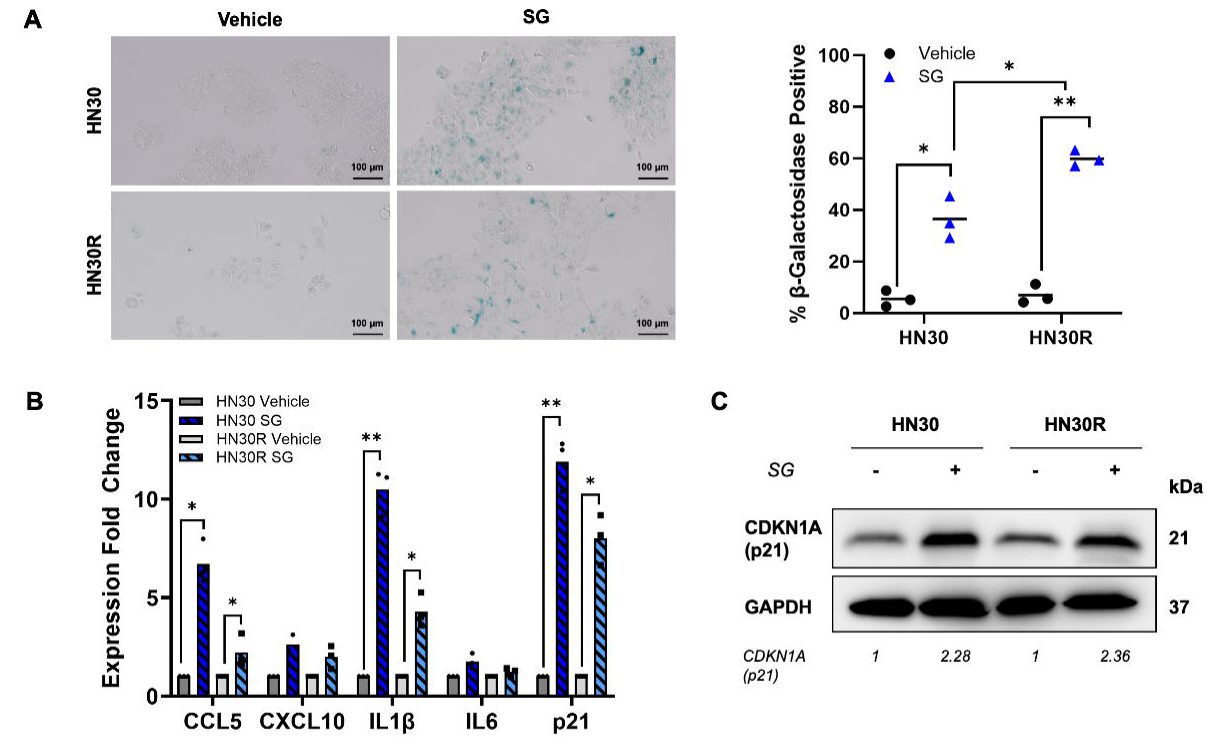
SG induces senescence in cisplatin-sensitive and -resistant HNSCC cells. **(A)** Left panel: HN30 and HN30R cells were treated with either 0.9% saline vehicle or 0.2 μg/mL SG for 72 hours and subjected to *in vitro* X-gal staining. Blue staining reflects β-galactosidase (β-gal) positive (senescent) cells. Right panel: Cells were stained with DDAOG following SG treatment and sorted via FACS to quantify senescence population via β-galactosidase activity. Each β-galactosidase positive population is representative of three biological replicates. **(B)** HN30 and HN30R cells were treated with either vehicle or 0.2 μg/mL SG for 72 hours and harvested for RNA isolation. RNA samples were then implemented for qPCR analysis with the indicated primers. Each value is representative of three technical replicates. Fold changes were calculated by considering the value of vehicle-treated HN30 or HN30R cells as 1. **(C)** HN30 and HN30R cells were treated as described in (**B**) and harvested for total cell lysates. Sixty micrograms of protein for each sample were loaded for western blot analysis via the indicated antibodies. Western blot data was analyzed via ImageJ quantification as described in Fig. 1C. \**p < 0.05, **p < 0.01*.

To further establish senescence induction via SG treatment in HN30 and HN30R cells, we utilized qPCR to assess the expression of representative SASPs, CCL5, CXCL10, IL6, and IL1β, as well as the cell cycle marker p21/CDKN1A (**Fig. 2B**). Both cell lines reflected upregulation of CCL5, CXCL10, IL6 and IL1β following SG treatment supporting senescence induction, though the level of SASP upregulation between the two cell lines differs for CCL5 and IL1β (**Fig. 2B**). Both RNA and protein levels of p21, an indicator of senescence, were significantly induced with SG treatment in both HN30 and HN30R cells (**Figs. 2B and 2C**). Together, these data support that SG induces senescence in both cisplatin-sensitive and - resistant HNSCC cells.

### The efficacy of SG-induced senescence is dependent on TROP2 expression and SN-38

TROP2 expression has been shown to dictate overall tumor response to SG treatment for multiple cancer types, including TNBC, ovarian cancer, and uterine carcinoma ^37^. Patients with these cancers who have higher levels of TROP2 expression have been found to respond more durably and display prolonged progression-free survival compared to those with lower TROP2 expression^37,38^. Though there are currently no studies examining TROP2 expression as a biomarker for HNSCC patient response, these studies suggest that TROP2 may be a necessary component of the delivery mechanism of SG and thus impact on its overall efficacy.

To assess the involvement of TROP-2 in the action of SG within HNSCC cells, we transduced HN30 cells with either a shControl or two shTROP2 constructs. We confirmed that both shTROP2 constructs significantly reduced TROP2 mRNA and protein levels (**Figs. 3A-B**). DDAOG FACS analysis revealed a significant decrease in senescence induction following SG treatment in HN30 cells expressing shTROP2 compared to shControl (**Fig. 3C**). These results support the anticipated outcome that the TROP2 protein plays a central role in the effectiveness of senescence induction by SG.

**Figure 3:**
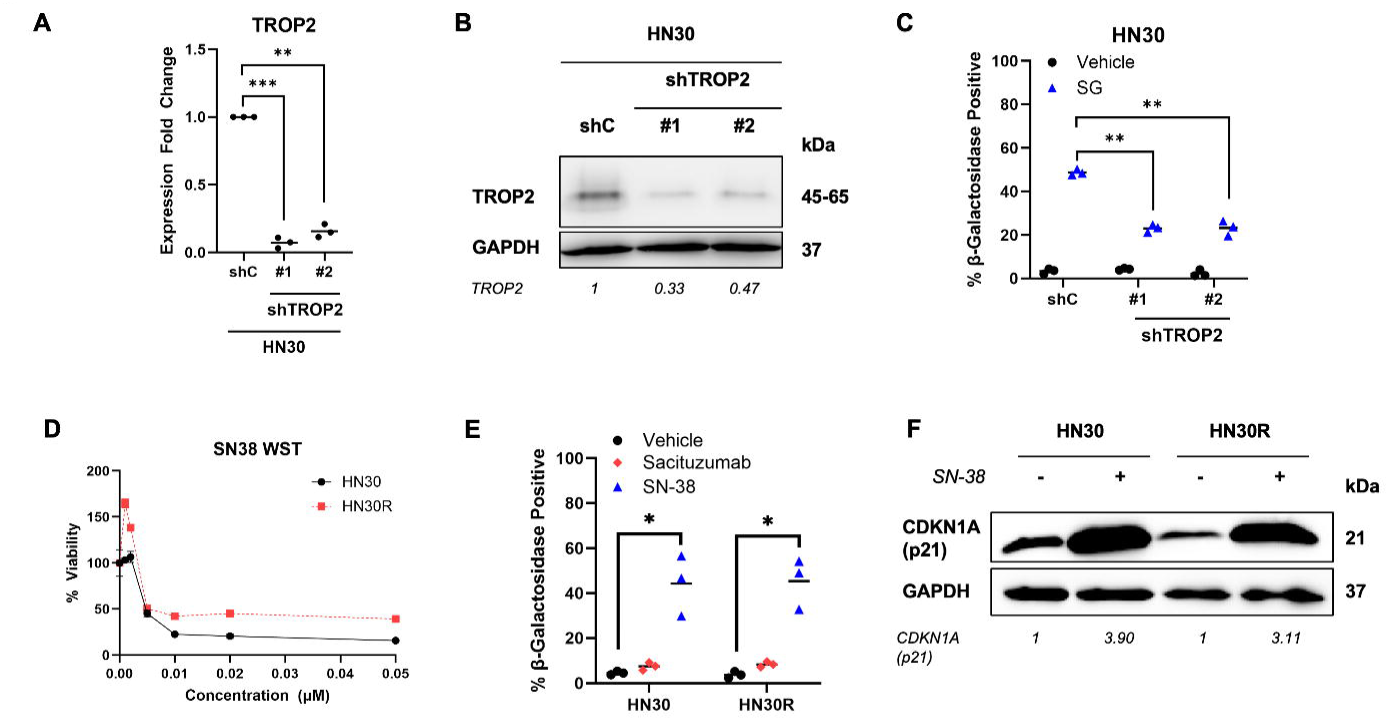
The efficacy of SG-induced senescence is dependent on TROP2 expression and SN-38. **(A)** HN30 cells were transfected with either a shControl (shC) or two different shTROP2 constructs (#1, #2). Clones from each transfectant were isolated and harvested for total RNA. RNA samples were then subjected to qPCR analysis for TROP2 expression; each value is indicative of three technical replicates. **(B)** Sixty micrograms of total cell lysates were loaded for western blot analysis for TROP2 expression. Densitometry was assessed via ImageJ quantification as described in Fig. 1C; fold change was calculated by comparing each sample to the TROP2 level of HN30 shC cells. **(C)** HN30 shC and shTROP2 expressing cells were treated with 0.2 μg/mL SG for 72 hours and subjected to β-gal positive cell quantitation via DDAOG FACS. **(D)** HN30 and HN30R cells were treated with 5 nM SN-38 for 72 hours and assessed for viability via WST-1 assay; each value is indicative of three biological replicates. **(E)** HN30 and HN30R cells were treated with either DMSO vehicle, 2.5 μg/mL bare sacituzumab antibody, or 5 nM SN-38 for 72 hours and subjected to DDAOG FACS to assess senescence population. All DDAOG FACS data are indicative of three biological replicates. **(F)** HN30 and HN30R cells were treated with 5 nM SN-38 and then harvested for total cell lysates. Sixty micrograms of protein were loaded for each sample for western blot analysis with the indicated antibodies. Densitometry was assessed via ImageJ quantification as described in Fig. 1C. \**p < 0.05, **p < 0.01, ***p<0.001*.

To confirm that SN-38, the SG warhead, contributes to SG-induced senescence (**Fig. 2**), we evaluated sensitivity to SN-38 in both cell lines with 72 hours of treatment. Both HN30 and HN30R cells displayed SN-38 IC_50_ concentrations of 5 nM (**Fig. 3D**). We then evaluated the extent of senescence induction following treatment with either vehicle, the naked sacituzumab monoclonal antibody, or the SN-38 warhead for 72 hours via DDAOG FACS analysis^39^. Both HN30 and HN30R cells showed significant induction of senescence, on the order of 45% senescence with SN-38 treatment while the bare sacituzumab antibody treatment did not induce a detectable level of senescence (**Fig. 3E**). Western blot analysis confirmed the induction of p21 following SN-38 treatment for both cell lines, as an additional indication of senescence (**Fig. 3F**). Taken together, these data indicate that SN-38 contributes to SG-mediated senescence induction in both cell lines once SG binds to TROP2 on the cell membrane.

### SG sensitizes cisplatin-sensitive and -resistant HNSCC cells to senolytic treatment

A recent phase II clinical basket trial of HNSCC patients found only a modest effect of SG as a single-agent over long-term treatment^21^. This limited effect may reflect SG-mediated senescence, although there were no efforts to detect senescence in patient tumors. Senolytics, drugs specifically targeting the senescent cell population, have been successful in a variety of solid tumors in combination with standard of care chemotherapies in preclinical studies^40^. We have demonstrated that ABT-263, a BH3-mimetic BCL-X_L_/BCL-2 dual inhibitor, sensitizes prostate cancer to androgen deprivation therapy, breast cancer to doxorubicin, head and neck cancer to cisplatin, and lung cancer to etoposide by eliminating the senescent cell population^28–30^. Consequently, we evaluated if a two-hit strategy of SG followed by senolytic treatment would be effective against both cisplatin-sensitive and -resistant HNSCC cells. We treated both HN30 and HN30R cells as well as Detroit-562 and Detroit-562R with SG for 72 hours followed by BH3-mimetics for 48 hours. We selected four compounds with specificity to three different pro-survival BCL-2 family proteins: ABT-263 (BCL-X_L_/BCL-2 inhibitor), S63845 (MCL-1 inhibitor), A-1155463 (BCL-X_L_ inhibitor), and ABT-199 (BCL-2 inhibitor). Crystal violet staining revealed that HN30 and HN30R cells as well as Detroit-562 and Detroit-562R cells have decreased overall viability when treated with either ABT-263 or A-1155463 following SG (**Fig. 4A**). These data suggest that senolytic activity observed here is dependent on BCL-X_L_ rather than on BCL-2 or MCL-1 as all four cell lines displayed unchanged cell viability when treated with either ABT-199 or S63835, respectively.

**Figure 4:**
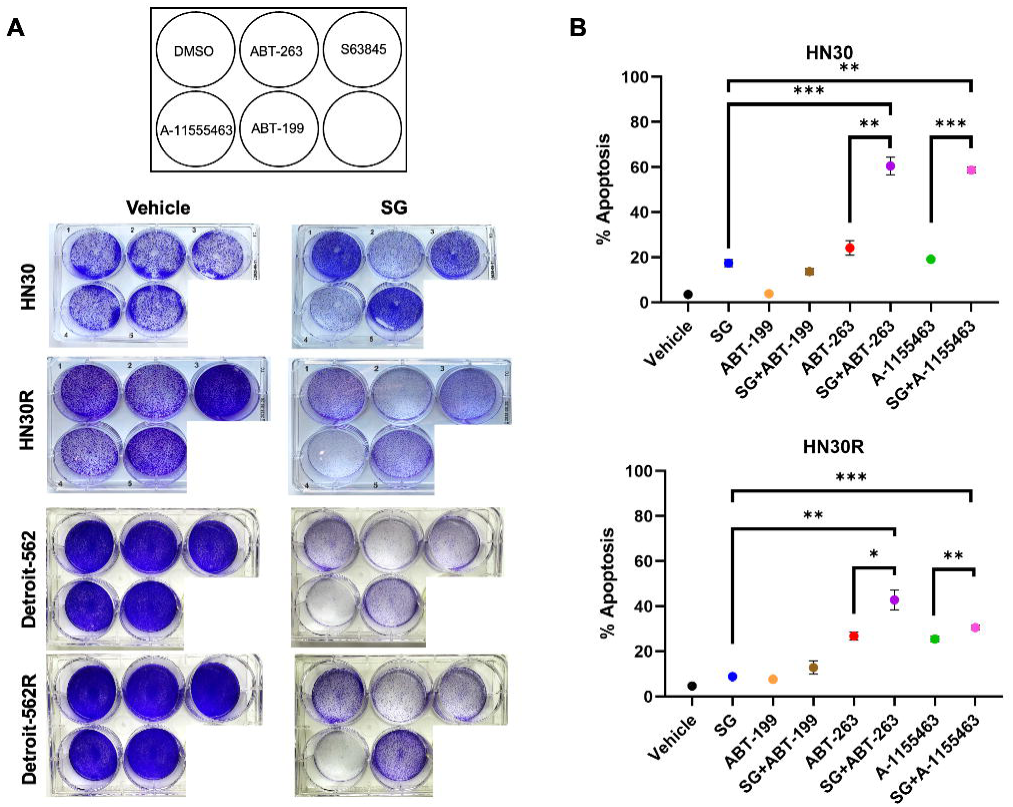
SG sensitizes cisplatin-sensitive and resistant HNSCC cells to BH3-mimetics. **(A)** HN30/HN30R and Detroit-562/Detroit-562R cells were treated with either 0.9% saline vehicle, 0.2 μg/mL SG (HN30/HN30R), or 1 μg/mL SG (Detroit-562/Detroit-562R) for 72 hours followed by treatment with either ABT-263 (1 μM), S63845 (0.1 μM), ABT-199 (1 μM), A-1155463 (1 μM), or DMSO for 48 hours. Cells were then stained with 0.05% Crystal Violet to assess for cell viability. **(B)** HN30 and HN30R cells were treated with 0.2 μg/mL SG for 72 hours followed by 1 μM of ABT-263, A-1155463, or ABT-199, respectively for 48 hours and assessed for apoptosis induction via Annexin-V/PI FACS; each treatment condition is representative for three biological replicates. *\*p < 0.05, **p < 0.01, ***p<0.001*.

We further assessed the promotion of apoptosis in response to the sequential treatment strategy of SG for 72 hours followed by either ABT-199, A-1155463, or ABT-263 for 48 hours via Annexin-V/PI FACS. Both HN30 and HN30R cells demonstrated a significant increase in apoptosis for SG combined with the BCL-X_L_-specific senolytics A-1155463 (HN30: 58.7%, HN30R: 30.5%) and ABT-263 (HN30: 60.5%, HN30R: 42.9%) compared to either SG, A-1155463, or ABT-263 alone (**Fig. 4B**). ABT-199 did not induce comparable levels of apoptosis either alone or in combination with SG in either HN30 (13.7%) or HN30R (12.8%) cells (**Fig. 4B**). Detroit-562 and Detroit-562R cells similarly displayed a significant increase in cell death when treated with SG for 72 hours followed by ABT-263 for 48 hours (Detroit-562: 54.76%; Detroit-562R: 68.7%) (**Supplemental Figure 2**). Together, these data support that SG-induced senescence followed by ABT-263 treatment is effective in sensitizing cisplatin-resistant HNSCC cells to cell death.

### Cell death induced by SG+ABT-263 is mediated by BAX and BAK

To examine the mechanism of action by which SG sensitizes cisplatin-sensitive and - resistant HNSCC cells to ABT-263, we assessed the expression of BCL-2 family proteins following treatment (**Fig. 5A**). Since the data presented in **Fig. 4** suggests that BCL-X_L_ plays a central role in senolytic activity, we focused on the expression of BCL-X_L_ and its pro-apoptotic interacting partners, BAX and BAK^41^. BCL-X_L_ expression was slightly induced with SG treatment alone in HN30 cells. However, HN30R did not demonstrate BCL-X_L_ upregulation with SG treatment. Pro-apoptotic BAK and BAX both demonstrated increased expression of their short forms following SG treatment, both alone and in combination with ABT-263. However, the intact forms of both BAK and BAX remained relatively unchanged between the treatment groups for both HN30 and HN30R cells. Cleaved-caspase-3 similarly displayed increased expression with SG treatment alone and further increased in combination with ABT-263, indicating increased cell death with these treatments and reflecting on the results in **Fig. 4B**.

**Figure 5:**
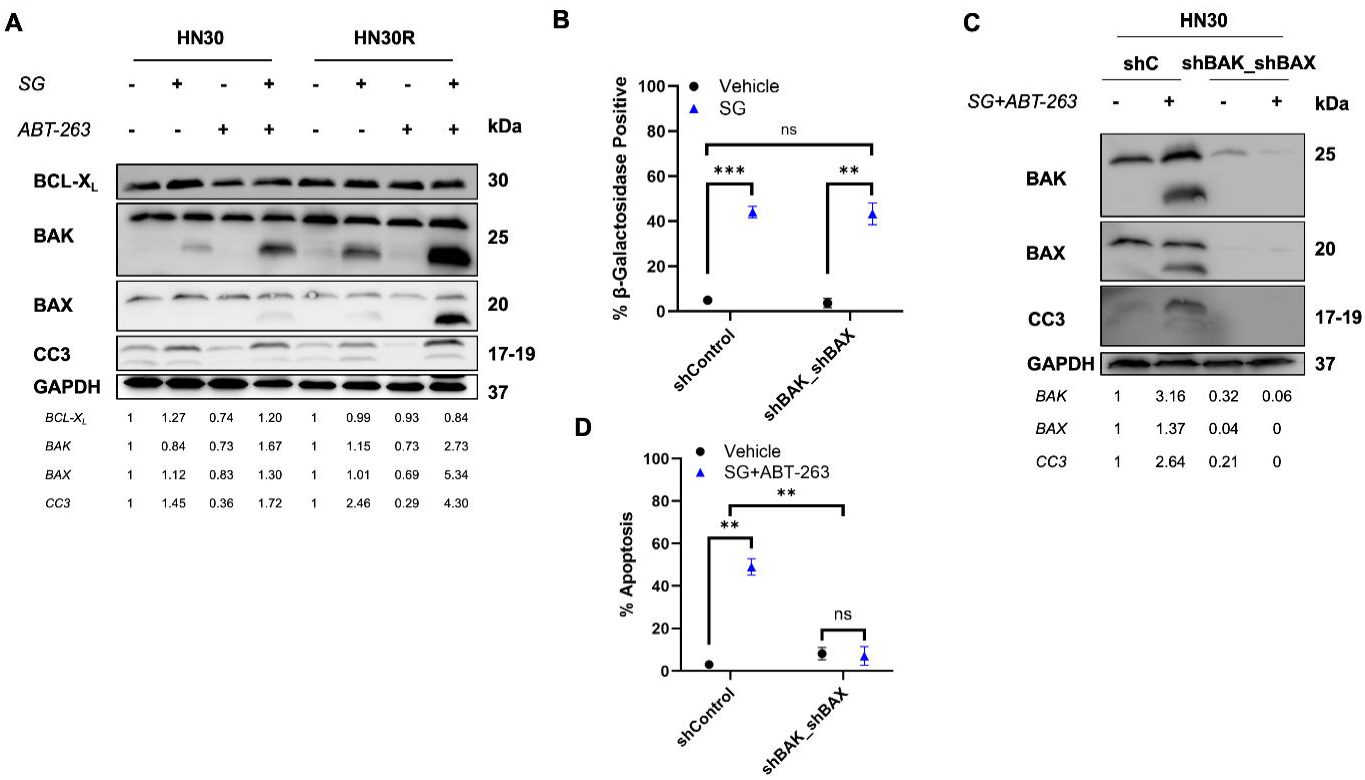
BAX and BAK are required for in SG + ABT-263-mediated senolysis. **(A)** HN30 and HN30R cells were treated with 0.2 μg/mL SG for 72 hours followed by 1 μM ABT-263 for 48 hours and then harvested for total cell lysates. Sixty micrograms of protein were loaded for each sample for western blot analysis with the indicated antibodies. Densitometry was assessed via ImageJ quantification as described in Fig. 1C. **(B)** HN30 shControl (shC) and shBAK/shBAX double-knockdown cells were treated with 0.2 μg/mL SG for 72 hours and evaluated for senescence induction via DDAOG FACS. **(C)** HN30 shC and shBAK/shBAX cells were treated with 0.2 μg/mL SG for 72 hours followed by 1 μM ABT-263 for 48 hours and harvested for total cell lysates for western blotting as described in **(A)**. **(D)** HN30 shC and shBAK/shBAX cells were treated as described in **(C)** and evaluated for apoptosis induction via Annexin-V/PI FACS. Each treatment condition is representative of three biological replicates. *\*\*p < 0.01, ***p<0.001*.

In order to further evaluate the role of BAK and BAX in apoptosis induction following SG and ABT-263 treatment, we established a double-knockdown BAK and BAX HN30 cell line that was treated with SG followed by ABT-263. Western blotting indicated continued BAK and BAX expression in HN30 shControl (shC) cells while shBAK/shBAX expressing HN30 cells displayed significantly reduced BAK and BAX expression (**Fig. 5C**). DDAOG β-galactosidase FACS revealed that knockdown of BAK and BAX did not impact senescence induction with SG treatment (**Fig. 5B**). However, cleaved-caspase-3 expression was undetectable in SG+ABT-263-treated HN30 shBAK/shBAX cells compared to shC expressing cells (**Fig. 5C**). Annexin-V/PI FACS indicated that BAK and BAX double-knockdown completely abrogated apoptosis induction with SG+ABT-263 treatment compared to shC in HN30 cells (**Fig. 5D**). Single-knockdown HN30 shBAK and shBAX cells also reflected decreased BAK and BAX expression, respectively, and displayed reduced expression of cleaved-caspase-3 and short form BAK and BAX with SG+ABT-263 treatment (**Supplemental Figure 3A**). In addition, single knockdowns of either BAK or BAX in HN30 cells reduced apoptosis induction about half compared to shC HN30 cells (∼60% in shC vs ∼30% in shBAK or shBAX) (**Supplemental Figure 3B**). These findings suggest that SG+ABT-263-mediated apoptotic cell death is dependent on both BAK and BAX expression.

### SG in combination with ABT-263 treatment limits tumor progression in vivo

Finally, we evaluated the durability of SG in combination with ABT-263 over time in cisplatin-resistant HN30R cells. HN30 and HN30R cells were treated with SG for 72 hours followed by ABT-263 for 48 hours and evaluated for live cell count at the indicated timepoints (**Fig. 6A**). SG treatment alone initially resulted in decreased cell viability; however, this viability was recovered after approximately 6 days following treatment (**Fig. 6A**). This is consistent with our previous studies displaying recovery from therapy-induced senescence in both cisplatin-sensitive and cisplatin-resistant HNSCC following treatment with chemotherapy^30,34^. Both HN30 and HN30R cells treated with SG and ABT-263 reflected prolonged decreased cell viability, however, HN30 cells completely recovered viability after approximately 9 days while HN30R recovered partial viability approximately 12 days after treatment (**Fig. 6A**). These data indicate that SG treatment alone is sufficient to induce senescence, though these cells ultimately recover proliferative capacity. Combination treatment with ABT-263, however, was effective in prolonging growth delay particularly in HN30R cells, nearly twice as long as SG treatment alone (**Fig. 6A**).

**Figure 6:**
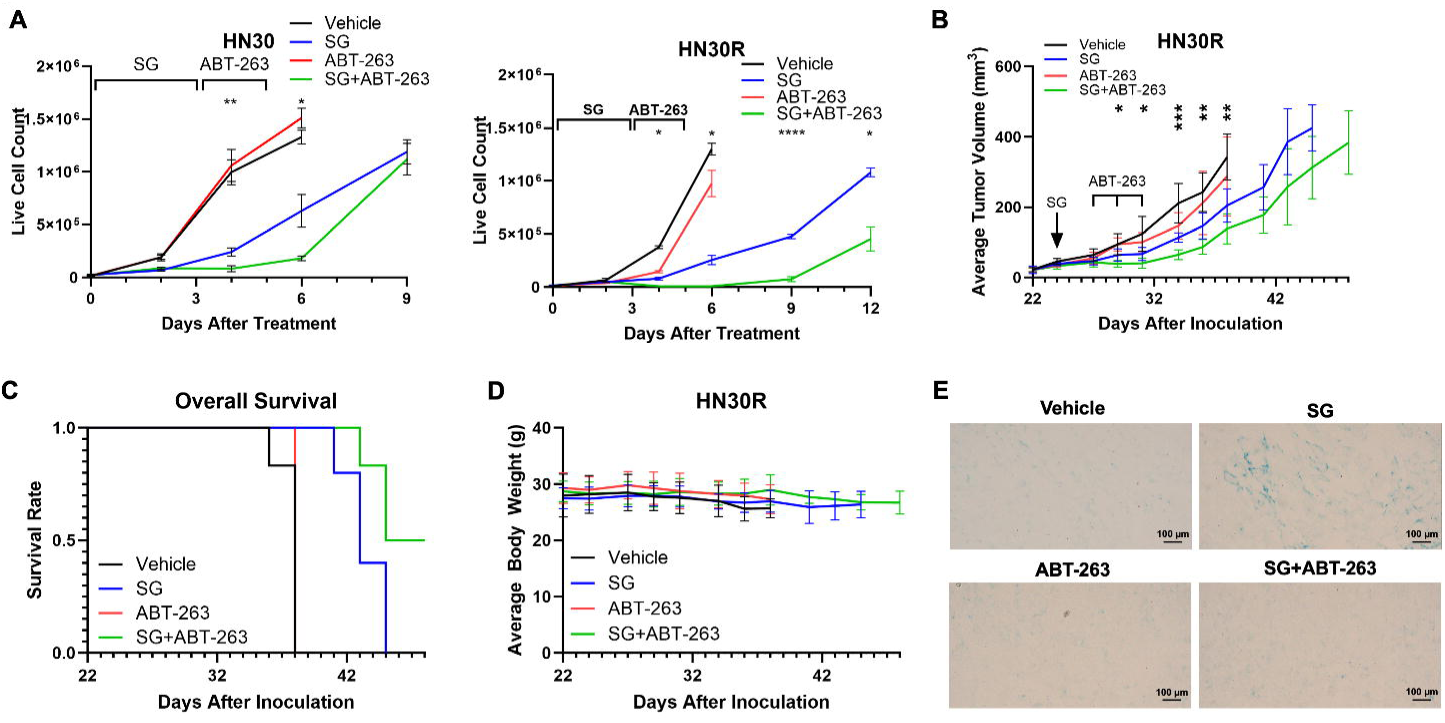
SG followed by ABT-263 treatment *in vivo* limits tumor progression in cisplatin-resistant HN30R tumors. **(A)** HN30 and HN30R cells were treated with 0.2 μg/mL SG for 72 hours followed by 1 μM ABT-263 for 48 hours and assessed for live cell count via trypan blue exclusion assay at the indicated timepoints. Each point represents the average of three biological replicates. Brackets display the treatment duration for SG and ABT-263. *p < 0.05, **p < 0.01, ***p<0.001, ****p<0.0001* (SG vs. SG+ABT-263). **(B)** HN30R (1x10^6^) cells were subcutaneously injected into the left cheek of NSG mice. Mice were randomized into either vehicle, SG, ABT-263, or SG+ABT-263 treatment groups (*n* = 6 per group) once tumors reached approximately 50 mm^3^ (Day 22). SG (0.25 mg/kg) was administered via intraperitoneal injection on Day 22 while ABT-263 (80 mg/kg) was given via oral gavage on Days 27, 29, and 31. Tumor volume was assessed via caliper measurement. \**p < 0.05, **p < 0.01, ***p<0.001* (vehicle vs. SG+ABT-263 treatment groups). **(C)** Overall survival over time was identified via Kaplan-Meier analysis. **(D)** Body weight was monitored throughout the study duration to assess drug toxicity. **(E)** Tumors were harvested from HN30R-inoculated NSG mice, stained via X-gal, and imaged to identify β-galactosidase positive cells (blue). The images are representative of three independent tumors in each treatment condition.

As this combination strategy was notably effective in delaying proliferative recovery in HN30R cells, we next examined the efficacy of this treatment *in vivo*. NSG mice were orthotopically inoculated with HN30R cells and monitored until their tumors reached ∼50 mm^3^ in volume. Mice were then randomized into four treatment groups (n = 6 per group): vehicle, SG-only, ABT-263-only, and SG followed by ABT-263 (SG+ABT-263). SG-only treated mice displayed reduced tumor progression compared to vehicle or ABT-263-only treated mice (**Fig. 6B**). Furthermore, mice sequentially treated with SG followed by ABT-263 displayed significantly delayed tumor growth over time compared to mice in other treatment groups (**Fig. 6B**). Specifically, SG+ABT-263 treated mice have a significantly lower tumor volume compared to vehicle at Day 29, 31, 34, 36, and 38 after inoculation. This is also consistent with the *in vitro* results displaying delayed proliferative recovery over time in HN30R cells treated with SG followed by ABT-263 (**Fig. 6A**). We also noted that SG-only and SG+ABT-263 treated mice displayed longer overall survival compared to vehicle or ABT-263 treated mice, with SG+ABT-263 mice reflecting the highest survival rate among the treatment groups (**Fig. 6C**). Body weight was not significantly affected by SG and/or ABT-263 treatment, indicating limited drug toxicity (**Fig. 6D**). These results were collectively replicated in an identical study (*n* = 8 per group), with the combination strategy of SG and ABT-263 significantly limiting tumor progression and improving overall survival with minimal effects on body weight (**Supplemental Figure 4**). We also confirmed that SG induces senescence *in vivo* via histochemical X-gal staining of tumor sections (**Fig. 6E**). Vehicle and ABT-263 only treated tumors displayed limited β-galactosidase positive cells while SG-only treated tumors displayed prominent β-galactosidase cell populations. However, SG+ABT-263 treated tumors displayed abrogated β-galactosidase positivity potentially as a result of ABT-263 promoting cell death in SG-induced senescent cell populations. Together, these data suggest that SG can induce senescence *in vivo* and may prove to be an effective strategy in combination with ABT-263 for limiting tumor growth to overcome cisplatin-resistance.

## Discussion

Antibody-drug conjugates (ADCs) have emerged as an innovative and efficient form of chemotherapy due to their ability to deliver their cytotoxic warhead to a targeted tumor cell population, thus circumventing off-target effects and limiting severe side effects. Sacituzumab govitecan (SG) has shown promising effects in patients with pretreated TNBC and hormone-receptor positive HER-2 negative breast cancer, increasing both overall and progression-free survival (PFS)^15,16^. Furthermore, SG has been shown to reduce overall tumor burden in platinum-resistant ovarian cancer and to decrease lesion number in urothelial cancer patients^18,42^. However, SG has had limited long-term success in pretreated HNSCC patients and has additionally displayed little effect as a single-agent^21^. This may potentially be attributed to senescence induction following SG treatment, preventing a durable, long-term response as the senescent populations eventually recover their proliferative capacity, leading to tumor relapse. Our previous studies support that both SG and its SN-38 warhead effectively induce senescence followed by proliferative recovery across a variety of solid tumors, including breast, prostate, lung, ovarian, and colorectal cancer^35^.

We show here that both cisplatin-sensitive HN30 and -resistant HN30R cells are sensitive to SG treatment; however, HN30R cells are significantly more responsive to SG. Our previous paper evaluating senolytic efficacy in HN30R cells following cisplatin treatment indicated that HN30R cells did not become senescent with cisplatin and thus failed to respond to the senolytic ABT-263 as a result of acquired cisplatin-resistance^34^. Yet, our current studies presented here display that SG effectively promotes senescence in HN30R cells and further primes these tumor cells for senolytic intervention. Notably, HN30R cells had a significantly higher level of senescence compared to HN30 parental cells following SG treatment. HN30R cell sensitivity to SG may also be a consequence of their increased levels of TROP2 expression compared to HN30 cells, suggesting that increased TROP2 expression may dictate SG sensitivity in HNSCC cells. In this context, we established that TROP2 expression is necessary for SG efficacy (**Fig. 3**), where decreased expression of TROP2 leads to decreased senescence induction following SG treatment. Additionally, we observed that SN-38 induces approximately equivalent levels of senescence in both HN30 and HN30R cells regardless of cisplatin-resistance (**Fig. 3E**), suggesting that SN-38 activity is not impacted by chemoresistance status. Studies evaluating digital pathology-based screening methodologies found that both pancreatic and lung cancer patients expressing high levels of TROP2 were the most responsive to treatment with TROP2-targeting therapies both in overall response rate and in PFS^43–45^. Both preclinical studies in esophageal adenocarcinoma and clinical studies in HER2-negative metastatic breast cancer have also indicated that TROP2 expression may impact overall sensitivity and PFS to SG treatment^46,47^. Thus, TROP2 expression may serve as an effective biomarker for stratifying HNSCC patients for SG treatment.

We observed that SG induces senescence in both cisplatin-sensitive and -resistant HSNCC cells (**Fig. 2**); however, both cell lines ultimately recovered their proliferative capacity following SG treatment alone (**Fig. 6A**). This potential recovery is delayed by exposure to the senolytic, ABT-263 in both cell lines. *In vivo* studies further supported the effectiveness of this strategy where HN30R-inoculated mice treated with SG followed by ABT-263 displayed reduced tumor progression and increased survival compared to mice treated with SG alone, indicating that ABT-263 may delay tumor relapse following SG-induced senescence (**Fig. 6B-C, Supplemental Fig. 4**). A recent study evaluating the efficacy of a TROP2-targeting ADC, sac-TMT, in NSCLC found that ADC treatment successfully limited the growth of drug-tolerant persister cells and significantly delayed tumor progression^48^. Our combination strategy of SG followed by ABT-263 may be an effective strategy for overcoming not only cisplatin-resistance but also SG resistance via targeting the senescent cell population.

Mechanistically, the combination of SG treatment followed by ABT-263 significantly upregulated apoptosis (**Fig. 4**). Crystal violet staining (**Fig. 4A**) confirmed that SG+ABT-263-mediated cell death is dependent on BCL-X_L._ Furthermore, double knockdown of both BAK and BAX abrogates apoptosis following SG+ABT-263 treatment without impacting senescence induction (**Figs. 5B-5D**), indicating that BAK and BAX predominantly regulate SG+ABT-263-mediated cell death independently of senescence induction. The short, presumably active forms of both BAK and BAX were highly expressed with this combination strategy, with moderate expression present with even SG treatment alone (**Fig. 5A**). Previous studies have shown that BAK and BAX may “auto” activate themselves, leading to increased homodimer formation and promoting mitochondrial permeabilization^49^. Furthermore, BAK has been shown to activate mitochondrial BAX, with increasing expression of its cleaved form^49,50^. Knockdown of BAK results in decreased expression of cleaved BAX following SG and ABT-263 treatment while BAX knockdown produces decreased cleaved BAK expression (**Supplemental Fig. 3A**). These data suggest that BAK and BAX may be inter-dependent to develop their cleaved, active forms.

Though ABT-263 is no longer being evaluated in clinical trials due to challenges with thrombocytopenia, it holds key mechanistic characteristics that are maintained by its later modified derivatives. DT2216, a VHL PROTAC BCL-X_L_ degrader, was recently evaluated in a phase I clinical trial in patients with relapsed solid tumors, including colorectal, ovarian, and pancreatic cancer^51,52^. Therefore ABT-263 may provide a proof-of-principle key mechanistic insight into the efficacy of alternative BCL-X_L_ inhibitors in combination with SG in the future.

This study focused on HPV-negative HNSCC as they comprise the significant majority of HNC cases and are dominated by advanced tumor progression and increased incidence of tumor relapse, typically requiring more aggressive treatment strategies. Studies evaluating TROP2 expression in HNSCC have indicated that there is no significant difference in TROP2 expression between HPV-negative and HPV-positive HNSCC tumors^53^, suggesting that both tumor subtypes may be responsive to SG. Future studies evaluating SG and ABT-263 efficacy will require analysis in HPV-positive cell lines in order to fully confirm the applicability of this combination in HNSCC.

These studies collectively indicate that a two-hit strategy of SG followed by ABT-263 treatment may be an effective method for suppressing cisplatin-resistant HNSCC tumors. Furthermore, SG may serve as an efficient and selective senescence inducer in tumor cell populations, allowing for more targeted delivery of senolytic therapeutics. Senolytics such as ABT-263 may be influential in mitigating acquired resistance to SG by eliminating senescent cell populations, thus prolonging the antitumor effects of this strategy.

## Acknowledgements

We thank Dr. Andrew Yeudall (Augusta University) for providing the HN30 cell line and Dr. Molly Bristol (VCU) for providing the N/Tert-1 keratinocyte lysates. We also thank the VCU Cancer Mouse Models Core and the VCU Flow Cytometry Core for their resources and experimental assistance.

## Author Contribution Statement

Conceptualization: N.L., D.A.G., H.H., Study design & consultation: N.L., D.A.G., H.H., J.K., methodology: N.L., B.H., data analysis: N.L., D.A.G., H.H., writing and draft preparation: N.L., D.A.G., H.H. All authors have read and agreed to the published version of the manuscript.

## Data Availability Statement

All original data and images are available within the manuscript.

## Ethics Statement

All animal studies were conducted in accordance with VCU Institutional Animal Care and Use Committee regulations.

---

**Supplementary Figure 1: SG induces senescence in parental Detroit-562 and cisplatin-resistant Detroit-562R cells.** Detroit-562 and Detroit-562R cells were treated with either 0.9% saline or 1 μg/mL SG for 72 hours, stained with DDAOG, and assessed via flow cytometry to quantify senescence induction via β-galactosidase activity. Each β-galactosidase positive population is representative of three biological replicates. *\*\*\*p<0.001, ****p<0.0001*.

**Supplementary Figure 2: SG sensitizes parental and cisplatin-resistant Detroit-562 cells to senolytic-induced cell death.** Detroit-562 and Detroit-562R cells were treated with 1 μg/mL SG for 72 hours followed by 1 μM of ABT-263 for 48 hours and assessed for apoptosis induction by Annexin-V/PI flow cytometry; each treatment condition is representative of three biological replicates. *\*\*p<0.01, ***p<0.001*.

**Supplementary Figure 3: Disruption of BAK or BAX individually is sufficient to reduce SG-mediated cell death following senolytic treatment. (A)** HN30 shControl (shC), shBAK, and shBAX cells were treated with 0.2 μg/mL SG for 72 hours followed by 1 μM ABT-263 for 48 hours and harvested for total cell lysates. Sixty micrograms of protein were loaded per sample for western blot analysis with the indicated antibodies. ImageJ was implemented for densitometric analysis as described in Fig. 1C. **(B)** HN30 shC, shBAK, and shBAX cells were treated as described in **(A)** and evaluated for apoptosis induction via Annexin-V/PI FACS; each condition is representative of three biological replicates. *\*\*p < 0.01*.

**Supplementary Figure 4: *In vivo* SG treatment limits tumor progression and improves overall survival in cisplatin-resistant HN30R tumors.** HN30R cells were subcutaneously injected into the left cheek of NSG mice. Mice were randomized into four groups (*n* = 8 per group) once their respective tumors reached approximately 50 mm^3^ (Day 13): vehicle (0.9% saline), SG (0.25 mg/kg administered via intraperitoneal injection), ABT-263 (80 mg/kg administered via oral-gavage), and SG+ABT-263. Mice received SG on Day 13 and ABT-263 treatment on Day 20, 22, and 24. **(A)** Tumor volume was measured via calipers at the indicated timepoints. **(B)** Kaplan-Meier analysis was implemented to examine overall survival. **(C)** Body weight was assessed at the indicated timepoints to evaluate toxicity.

## References

1 Florea A-M, Büsselberg D. Cisplatin as an Anti-Tumor Drug: Cellular Mechanisms of Activity, Drug Resistance and Induced Side Effects. Cancers 2011; 3: 1351–1371.

2 Johnson DE, Burtness B, Leemans CR, Lui VWY, Bauman JE, Grandis JR. Head and neck squamous cell carcinoma. Nat Rev Dis Primers 2020; 6: 92.

3 Saâd MA, Taleb I, El Omri S, Traoré H, Chahbounia I, Elm’hadi C et al. Comparing efficacy and safety of weekly vs. triweekly cisplatin concurrently with radiotherapy for locally advanced head and neck cancer: Systematic review and meta-analysis. Oral Oncology Reports 2025; 14: 100738.

4 Dasari S, Bernard Tchounwou P. Cisplatin in cancer therapy: Molecular mechanisms of action. European Journal of Pharmacology 2014; 740: 364–378.

5 Demaria M, O’Leary MN, Chang J, Shao L, Liu S, Alimirah F et al. Cellular Senescence Promotes Adverse Effects of Chemotherapy and Cancer Relapse. Cancer Discovery 2017; 7: 165–176.

6 Jeiroshi A, Deng J, Xu Z, Comandatore A, Xu G, Glaviano A et al. Navigating the paradox of senescence and chemoresistance in pancreatic cancer. Seminars in Cancer Biology 2025; 114: 60–72.

7 Khongorzul P, Ling CJ, Khan FU, Ihsan AU, Zhang J. Antibody–Drug Conjugates: A Comprehensive Review. Molecular Cancer Research 2020; 18: 3–19.

8 Park JC, Shin D. Current Landscape of Antibody-Drug Conjugate Development in Head and Neck Cancer. JCO Precis Oncol 2024; 8: e2400179.

9 Tsuchikama K, Anami Y, Ha SYY, Yamazaki CM. Exploring the next generation of antibody– drug conjugates. Nat Rev Clin Oncol 2024; 21: 203–223.

10 Tang G, Tang Q, Jia L, Xia S, Li J, Chen Y et al. High expression of TROP2 is correlated with poor prognosis of oral squamous cell carcinoma. Pathology - Research and Practice 2018; 214: 1606–1612.

11 Mas L, Cros J, Svrcek M, Van Laethem JL, Emile JF, Rebours V et al. Trop-2 is a ubiquitous and promising target in pancreatic adenocarcinoma. Clinics and Research in Hepatology and Gastroenterology 2023; 47: 102108.

12 Bardia A, Mayer IA, Vahdat LT, Tolaney SM, Isakoff SJ, Diamond JR et al. Sacituzumab Govitecan-hziy in Refractory Metastatic Triple-Negative Breast Cancer. N Engl J Med 2019; 380: 741–751.

13 Syed YY. Sacituzumab Govitecan: First Approval. Drugs 2020; 80: 1019–1025.

14 Zhang Y, Chen J, Wang X, Wang H, Chen X, Hong J et al. Efficacy and safety of Sacituzumab govitecan in solid tumors: a systematic review and meta-analysis. Front Oncol 2025; 15: 1624386.

15 Bardia A, Hurvitz SA, Tolaney SM, Loirat D, Punie K, Oliveira M et al. Sacituzumab Govitecan in Metastatic Triple-Negative Breast Cancer. N Engl J Med 2021; 384: 1529– 1541.

16 Rugo HS, Bardia A, Marmé F, Cortes J, Schmid P, Loirat D et al. Sacituzumab Govitecan in Hormone Receptor-Positive/Human Epidermal Growth Factor Receptor 2-Negative Metastatic Breast Cancer. J Clin Oncol 2022; 40: 3365–3376.

17 Faltas B, Goldenberg DM, Ocean AJ, Govindan SV, Wilhelm F, Sharkey RM et al. Sacituzumab Govitecan, a Novel Antibody–Drug Conjugate, in Patients With Metastatic Platinum-Resistant Urothelial Carcinoma. Clinical Genitourinary Cancer 2016; 14: e75–e79.

18 Greenman M, Bellone S, Demirkiran C, Max Philipp Hartwich T, Santin AD. Sacituzumab govitecan in heavily pretreated, platinum-resistant high grade serous ovarian cancer. Gynecologic Oncology Reports 2024; 54: 101459.

19 Okajima D, Yasuda S, Maejima T, Karibe T, Sakurai K, Aida T et al. Datopotamab Deruxtecan, a Novel TROP2-directed Antibody-drug Conjugate, Demonstrates Potent Antitumor Activity by Efficient Drug Delivery to Tumor Cells. Mol Cancer Ther 2021; 20: 2329–2340.

20 Sands J, Ahn M-J, Lisberg A, Cho BC, Blumenschein G, Shum E et al. Datopotamab Deruxtecan in Advanced or Metastatic Non-Small Cell Lung Cancer With Actionable Genomic Alterations: Results From the Phase II TROPION-Lung05 Study. J Clin Oncol 2025; 43: 1254–1265.

21 Michel L, Jimeno A, Sukari A, Beck JT, Chiu J, Ahern E et al. Sacituzumab Govitecan in Patients with Relapsed/Refractory Advanced Head and Neck Squamous Cell Carcinoma: Results from the Phase II TROPiCS-03 Basket Study. Clin Cancer Res 2025; 31: 832–838.

22 Park SS, Choi YW, Kim J-H, Kim HS, Park TJ. Senescent tumor cells: an overlooked adversary in the battle against cancer. Exp Mol Med 2021; 53: 1834–1841.

23 Xiao S, Qin D, Hou X, Tian L, Yu Y, Zhang R et al. Cellular senescence: a double-edged sword in cancer therapy. Front Oncol 2023; 13: 1189015.

24 Coppé J-P, Desprez P-Y, Krtolica A, Campisi J. The Senescence-Associated Secretory Phenotype: The Dark Side of Tumor Suppression. Annu Rev Pathol Mech Dis 2010; 5: 99– 118.

25 Nelson G, Wordsworth J, Wang C, Jurk D, Lawless C, Martin-Ruiz C et al. A senescent cell bystander effect: senescence-induced senescence. Aging Cell 2012; 11: 345–349.

26 Chen Y-F, Xu Y-Y, Shao Z-M, Yu K-D. Resistance to antibody-drug conjugates in breast cancer: mechanisms and solutions. Cancer Commun (Lond) 2023; 43: 297–337.

27 Saleh T, Greenberg EF, Faber AC, Harada H, Gewirtz DA. A Critical Appraisal of the Utility of Targeting Therapy-Induced Senescence for Cancer Treatment. Cancer Res 2025; 85: 1755– 1768.

28 Saleh T, Carpenter VJ, Tyutyunyk-Massey L, Murray G, Leverson JD, Souers AJ et al. Clearance of therapy-induced senescent tumor cells by the senolytic ABT-263 via interference with BCL-X _L_ –BAX interaction. Mol Oncol 2020; 14: 2504–2519.

29 Carpenter VJ, Patel BB, Autorino R, Smith SC, Gewirtz DA, Saleh T. Senescence and castration resistance in prostate cancer: A review of experimental evidence and clinical implications. Biochimica et Biophysica Acta (BBA) - Reviews on Cancer 2020; 1874: 188424.

30 Ahmadinejad F, Bos T, Hu B, Britt E, Koblinski J, Souers AJ et al. Senolytic-Mediated Elimination of Head and Neck Tumor Cells Induced Into Senescence by Cisplatin. Mol Pharmacol 2022; 101: 168–180.

31 Flor A, Pagacz J, Thompson D, Kron S. Far-red Fluorescent Senescence-associated β-Galactosidase Probe for Identification and Enrichment of Senescent Tumor Cells by Flow Cytometry. J Vis Exp 2022. doi:10.3791/64176.

32 James CD, Lewis RL, Fakunmoju AL, Witt AJ, Youssef AH, Wang X et al. Fibroblast stromal support model for predicting human papillomavirus-associated cancer drug responses. J Virol 2024; 98: e01024–24.

33 Oweida AJ, Bhatia S, Van Court B, Darragh L, Serkova N, Karam SD. Intramucosal Inoculation of Squamous Cell Carcinoma Cells in Mice for Tumor Immune Profiling and Treatment Response Assessment. J Vis Exp 2019. doi:10.3791/59195.

34 Luffman N, Ahmadinejad F, Finnegan RM, Raymond M, Gewirtz DA, Harada H. Effectiveness of PROTAC BET Degraders in Combating Cisplatin Resistance in Head and Neck Cancer Cells. IJMS 2025; 26: 6185.

35 Chakraborty E, Sinanian M, Rahman A, Gewirtz DA. Promotion of Tumor Cell Senescence by the Antibody–Drug Conjugate, Sacituzumab Govitecan. Molecular Pharmacology 2026; : 100142.

36 Debacq-Chainiaux F, Erusalimsky JD, Campisi J, Toussaint O. Protocols to detect senescence-associated beta-galactosidase (SA-βgal) activity, a biomarker of senescent cells in culture and in vivo. Nat Protoc 2009; 4: 1798–1806.

37 Qiu S, Zhang J, Wang Z, Lan H, Hou J, Zhang N et al. Targeting Trop-2 in cancer: Recent research progress and clinical application. Biochimica et Biophysica Acta (BBA) - Reviews on Cancer 2023; 1878: 188902.

38 Yao L, Chen J, Ma W. Decoding TROP2 in breast cancer: significance, clinical implications, and therapeutic advancements. Front Oncol 2023; 13: 1292211.

39 Nair JR, Huang T-T, Sunkara A, Pruitt MR, Ibanez KR, Chiang C-Y et al. Distinct effects of sacituzumab govitecan and berzosertib on DNA damage response in ovarian cancer. iScience 2024; 27: 111283.

40 Carpenter VJ, Saleh T, Gewirtz DA. Senolytics for Cancer Therapy: Is All That Glitters Really Gold? Cancers (Basel) 2021; 13: 723.

41 Shamas-Din A, Kale J, Leber B, Andrews DW. Mechanisms of action of Bcl-2 family proteins. Cold Spring Harb Perspect Biol 2013; 5: a008714.

42 Powles T, Tagawa S, Vulsteke C, Gross-Goupil M, Park SH, Necchi A et al. Sacituzumab govitecan in advanced urothelial carcinoma: TROPiCS-04, a phase III randomized trial. Ann Oncol 2025; 36: 561–571.

43 Ye L, Chen H, Wu D. TROP2-targeted molecular imaging: a promising tool for precision oncology. Am J Nucl Med Mol Imaging 2025; 15: 109–123.

44 Ahn M-J, Tanaka K, Paz-Ares L, Cornelissen R, Girard N, Pons-Tostivint E et al. Datopotamab Deruxtecan Versus Docetaxel for Previously Treated Advanced or Metastatic Non-Small Cell Lung Cancer: The Randomized, Open-Label Phase III TROPION-Lung01 Study. J Clin Oncol 2025; 43: 260–272.

45 Novel computational pathology-based TROP2 biomarker for datopotamab deruxtecan was predictive of clinical outcomes in patients with non-small cell lung cancer in TROPION-Lung01 Phase III trial. 2024.https://www.astrazeneca.com/content/astraz/media-centre/press-releases/2024/novel-computational-pathology-based-trop2-biomarker-for-dato-dxd-was-predictive-of-clinical-outcomes-in-patients-with-nsclc-in-tropion-lung01-phase-iii-trial.html (accessed 31 Mar2026).

46 Hoppe S, Meder L, Gebauer F, Ullrich RT, Zander T, Hillmer AM et al. Trophoblast Cell Surface Antigen 2 (TROP2) as a Predictive Bio-Marker for the Therapeutic Efficacy of Sacituzumab Govitecan in Adenocarcinoma of the Esophagus. Cancers (Basel) 2022; 14: 4789.

47 LeVee A, Wong M, Ruel N, Schmolze D, Han M, Mortimer J et al. Trop-2 expression as a biomarker of response to sacituzumab govitecan in patients with HER2-negative metastatic breast cancer: A pilot study. Cancer Treat Res Commun 2025; 44: 100954.

48 Liao J, Yao W, Yu Y, Li A, Huang J, Li X et al. Targeting TROP2 in drug-tolerant persister cells delays EGFR tyrosine kinase inhibitor resistance in non-small-cell lung cancer. Cancer Cell 2026; 44: 1401–1417.e9.

49 Iyer S, Uren RT, Dengler MA, Shi MX, Uno E, Adams JM et al. Robust autoactivation for apoptosis by BAK but not BAX highlights BAK as an important therapeutic target. Cell Death Dis 2020; 11: 268.

50 Singh G, Guibao CD, Seetharaman J, Aggarwal A, Grace CR, McNamara DE et al. Structural basis of BAK activation in mitochondrial apoptosis initiation. Nat Commun 2022; 13: 250.

51 Khan S, Zhang X, Lv D, Zhang Q, He Y, Zhang P et al. A selective BCL-XL PROTAC degrader achieves safe and potent antitumor activity. Nat Med 2019; 25: 1938–1947.

52 Mahadevan D, Barve M, Mahalingam D, Parekh J, Kurman M, Strauss J et al. First in human phase 1 study of DT2216, a selective BCL-xL degrader, in patients with relapsed/refractory solid malignancies. J Hematol Oncol 2025; 18: 98.

53 Klasen C, Eckel HNC, Wuerdemann N, Böckelmann J, Knipper K, Suchan M et al. Expression patterns of TROP2 and Nectin-4 in oropharyngeal squamous cell carcinoma in relation to HPV status: potential biomarkers for targeted therapy. Ther Adv Med Oncol 2025; 17: 17588359251361877.

