## Supplemental Data for "Sacituzumab Govitecan as an Effective Strategy for Sensitizing Chemoresistant HNSCC Cells to Senolytic Intervention"

### Slide 1
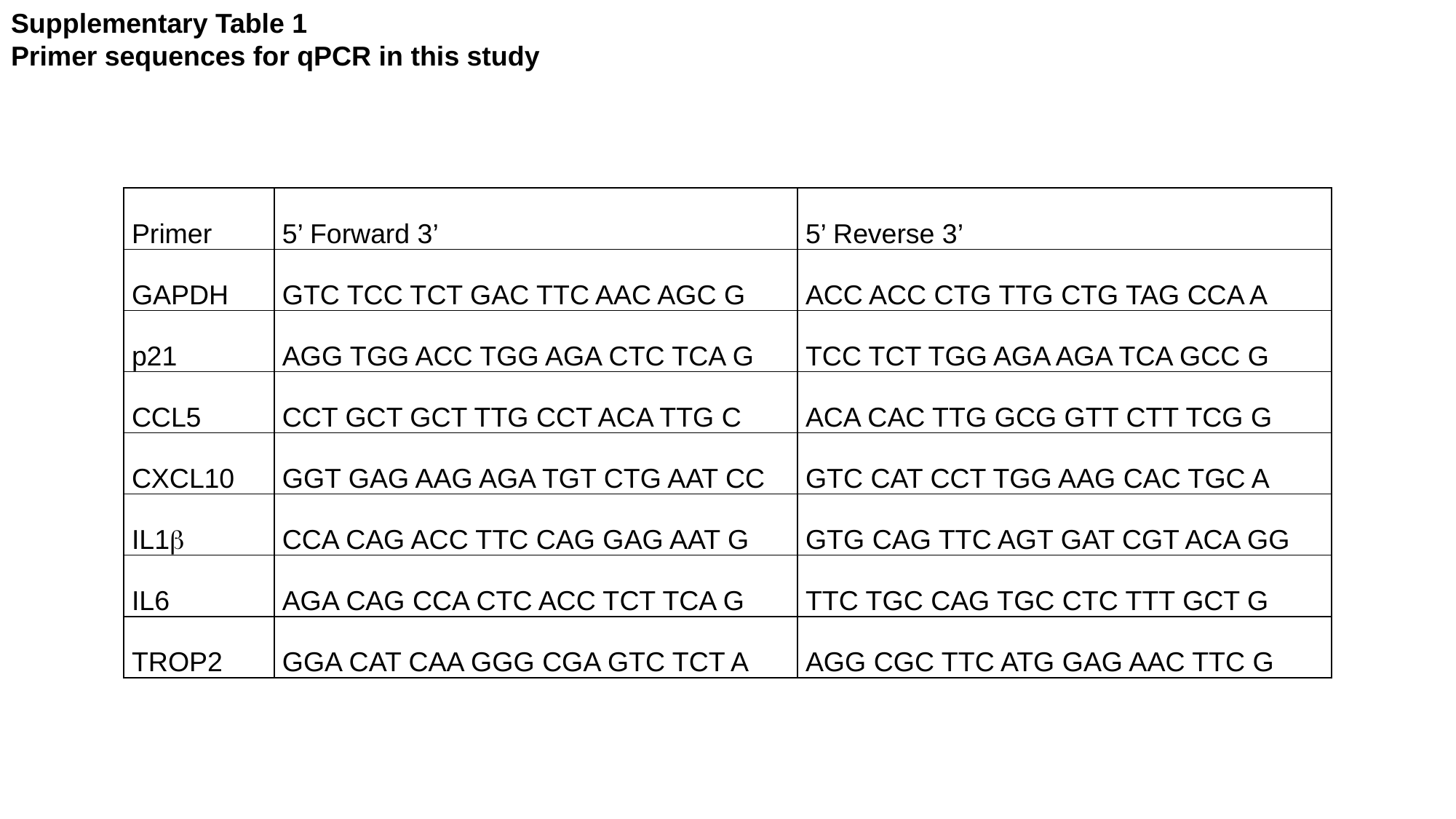

Supplementary Table 1
Primer sequences for qPCR in this study
| Primer | 5’ Forward 3’ | 5’ Reverse 3’ |
| --- | --- | --- |
| GAPDH | GTC TCC TCT GAC TTC AAC AGC G | ACC ACC CTG TTG CTG TAG CCA A |
| p21 | AGG TGG ACC TGG AGA CTC TCA G | TCC TCT TGG AGA AGA TCA GCC G |
| CCL5 | CCT GCT GCT TTG CCT ACA TTG C | ACA CAC TTG GCG GTT CTT TCG G |
| CXCL10 | GGT GAG AAG AGA TGT CTG AAT CC | GTC CAT CCT TGG AAG CAC TGC A |
| IL1β | CCA CAG ACC TTC CAG GAG AAT G | GTG CAG TTC AGT GAT CGT ACA GG |
| IL6 | AGA CAG CCA CTC ACC TCT TCA G | TTC TGC CAG TGC CTC TTT GCT G |
| TROP2 | GGA CAT CAA GGG CGA GTC TCT A | AGG CGC TTC ATG GAG AAC TTC G |

### Slide 2
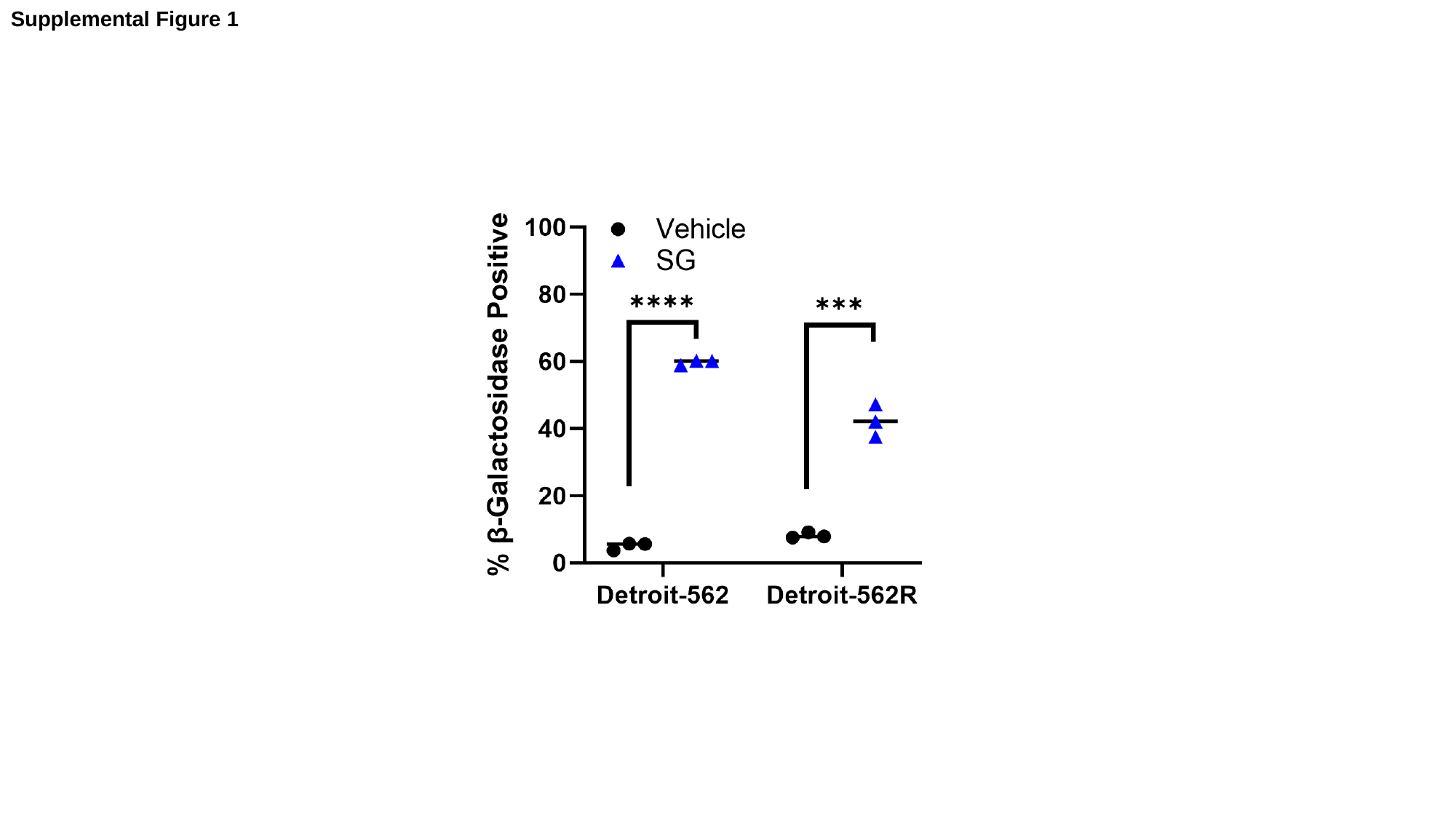

Supplemental Figure 1

### Slide 3
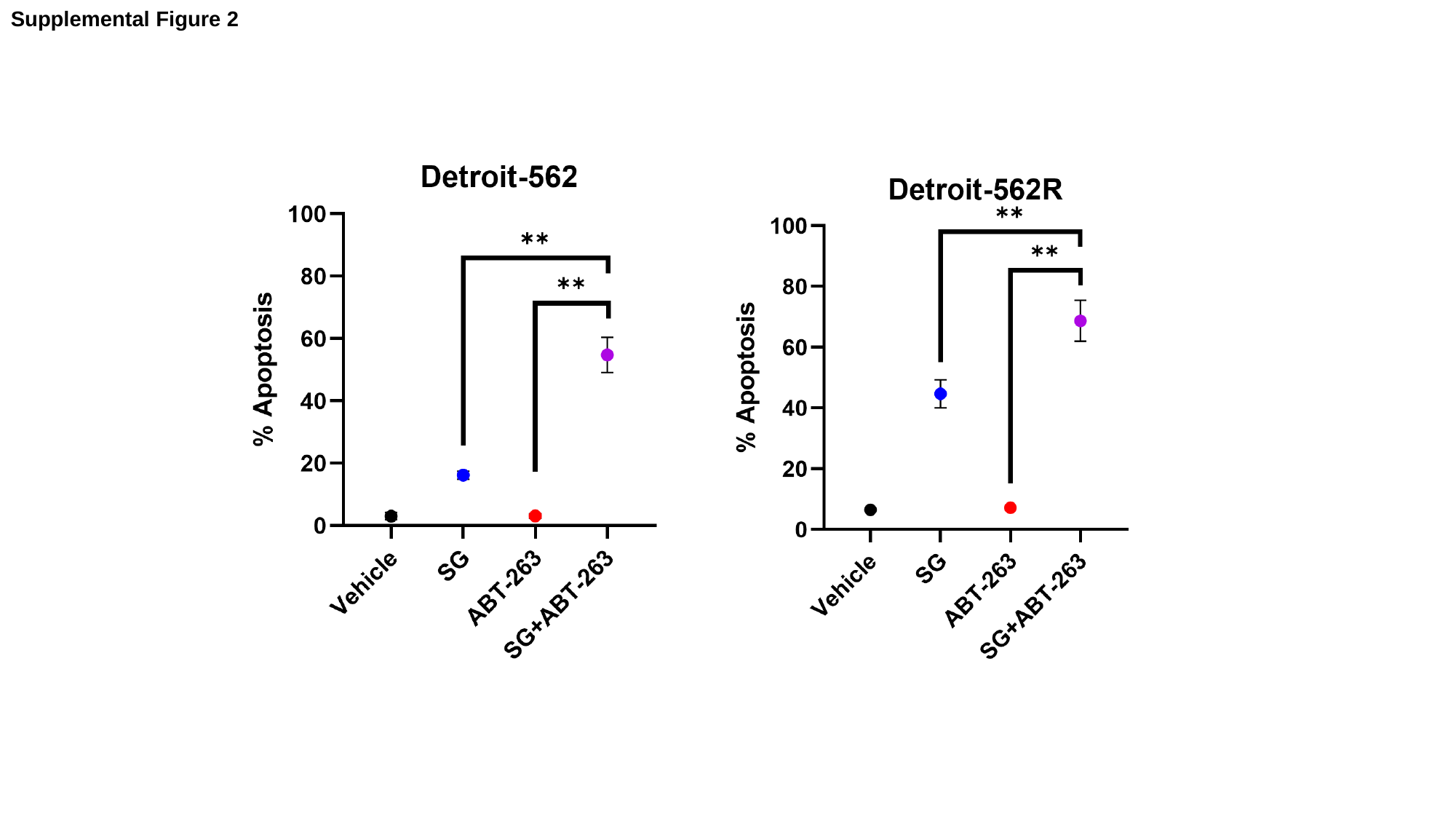

Supplemental Figure 2

### Slide 4
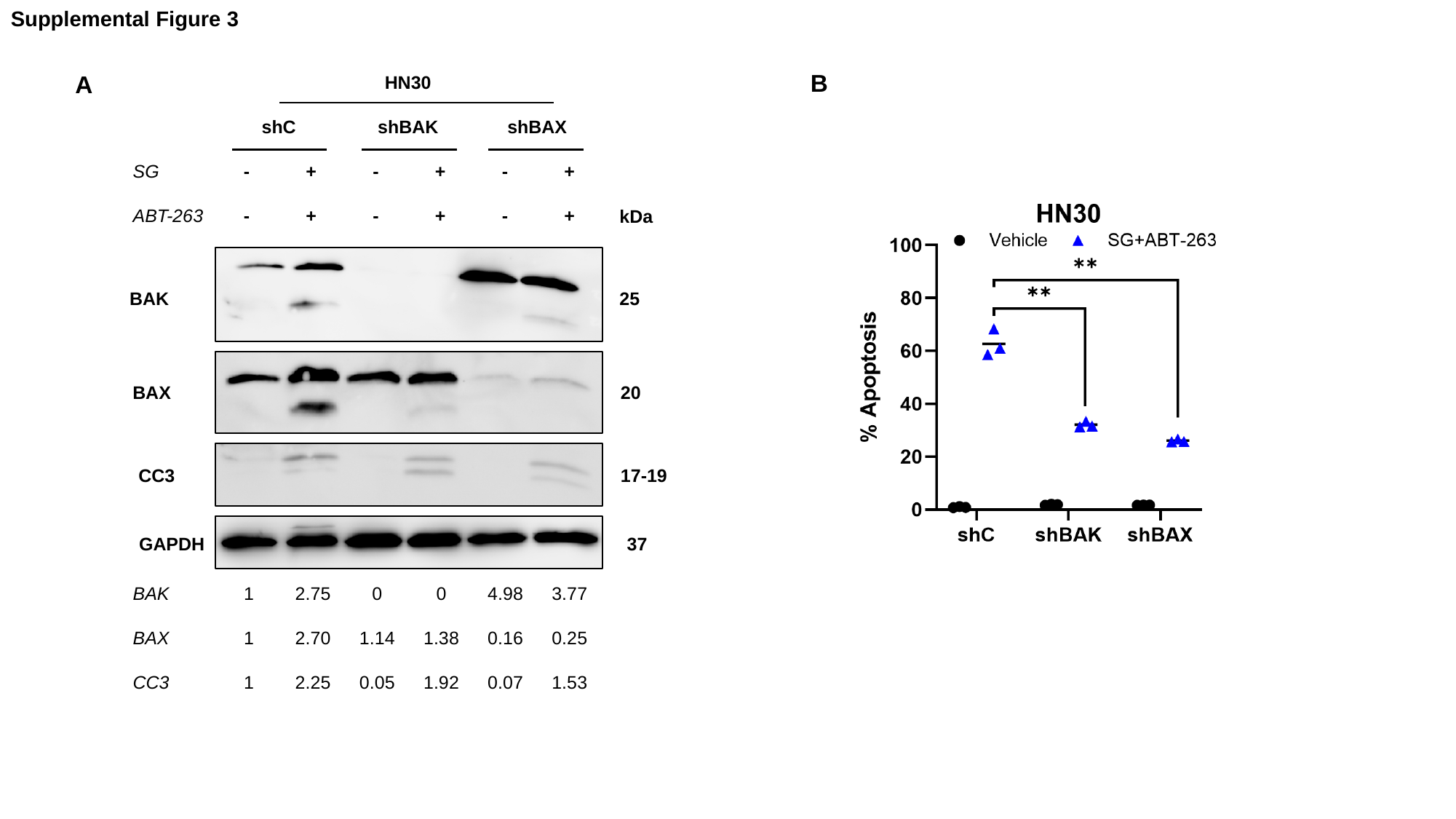

Supplemental Figure 3
B
A
| | HN30 | | | | | |
| --- | --- | --- | --- | --- | --- | --- |
| | shC | | shBAK | | shBAX | |
| SG | - | + | - | + | - | + |
| ABT-263 | - | + | - | + | - | + |
kDa
25
BAK
20
BAX
CC3
17-19
GAPDH
37
| BAK | 1 | 2.75 | 0 | 0 | 4.98 | 3.77 |
| --- | --- | --- | --- | --- | --- | --- |
| BAX | 1 | 2.70 | 1.14 | 1.38 | 0.16 | 0.25 |
| CC3 | 1 | 2.25 | 0.05 | 1.92 | 0.07 | 1.53 |

### Slide 5
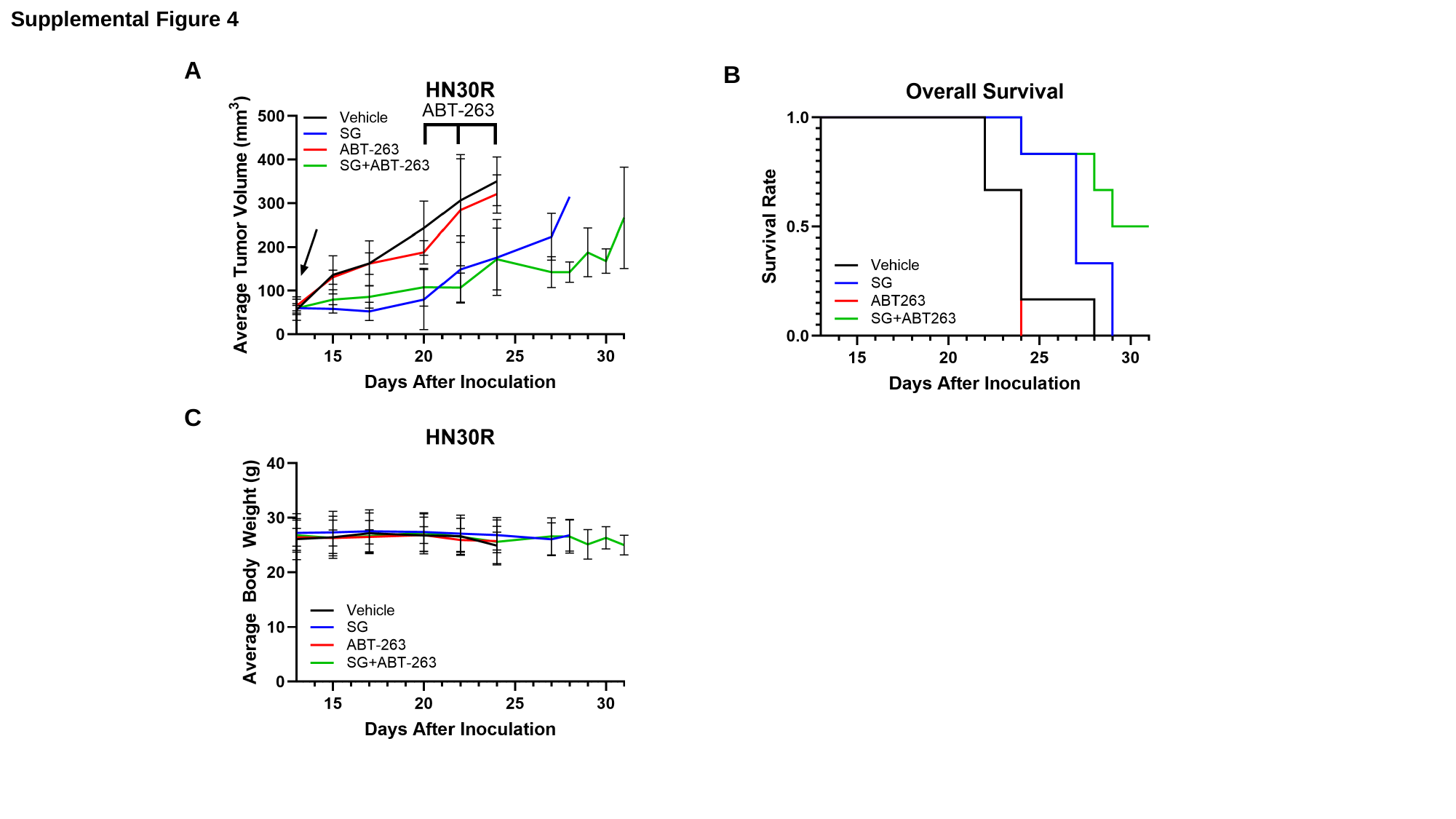

Supplemental Figure 4
A
B
C

### Slide 6
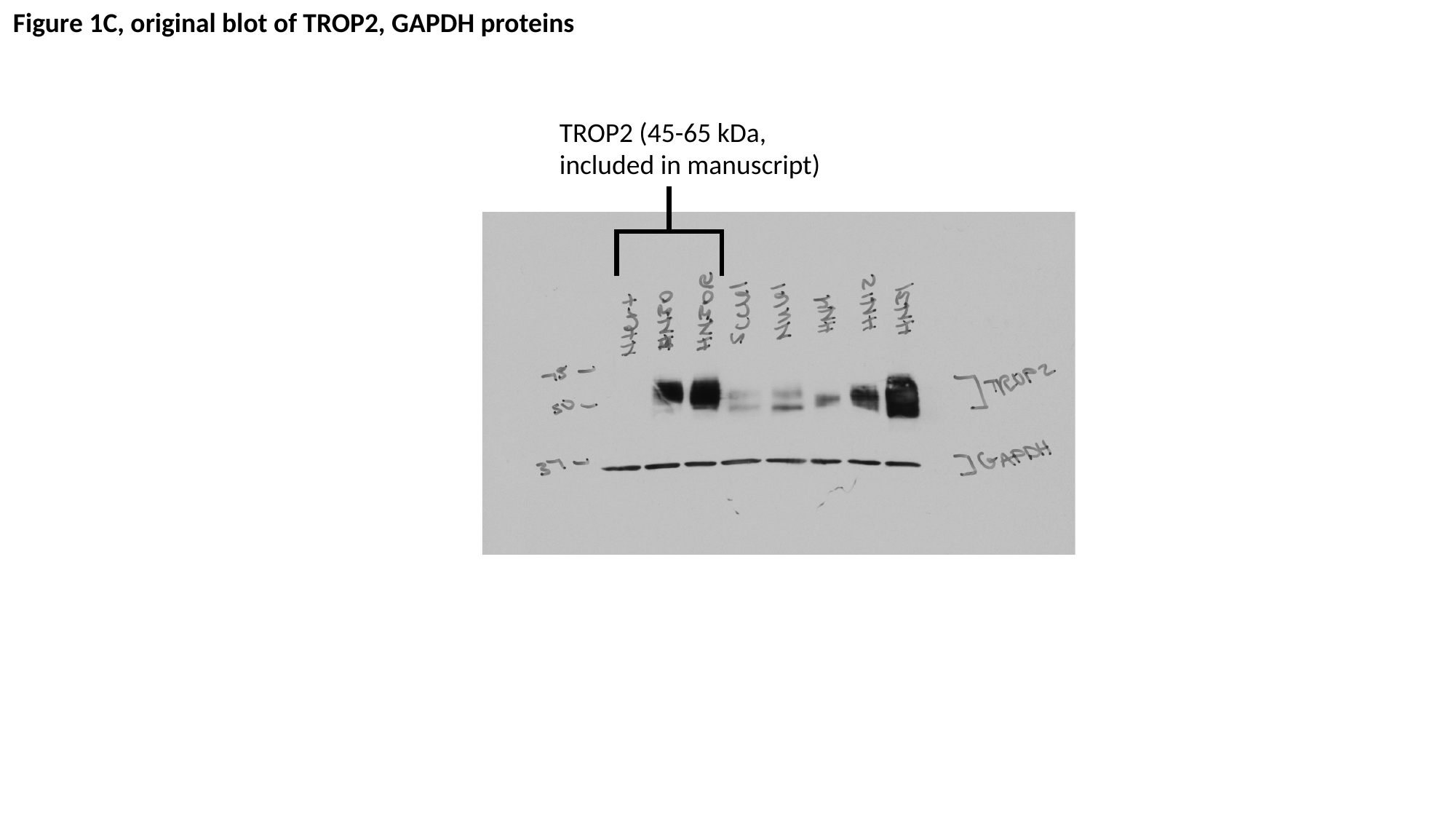

Figure 1C, original blot of TROP2, GAPDH proteins
TROP2 (45-65 kDa, included in manuscript)

### Slide 7
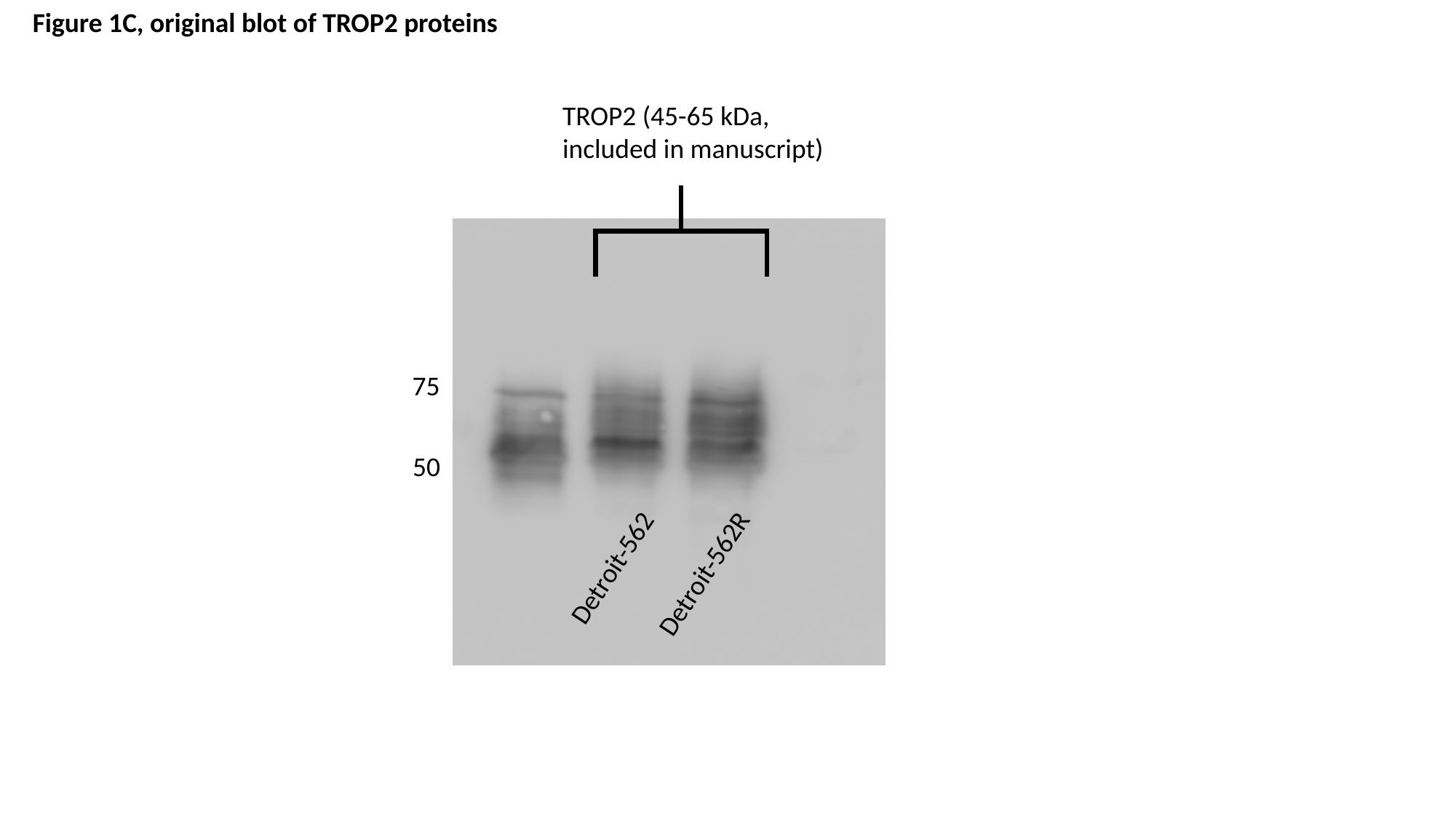

Figure 1C, original blot of TROP2 proteins
TROP2 (45-65 kDa, included in manuscript)
75
50
Detroit-562
Detroit-562R

### Slide 8
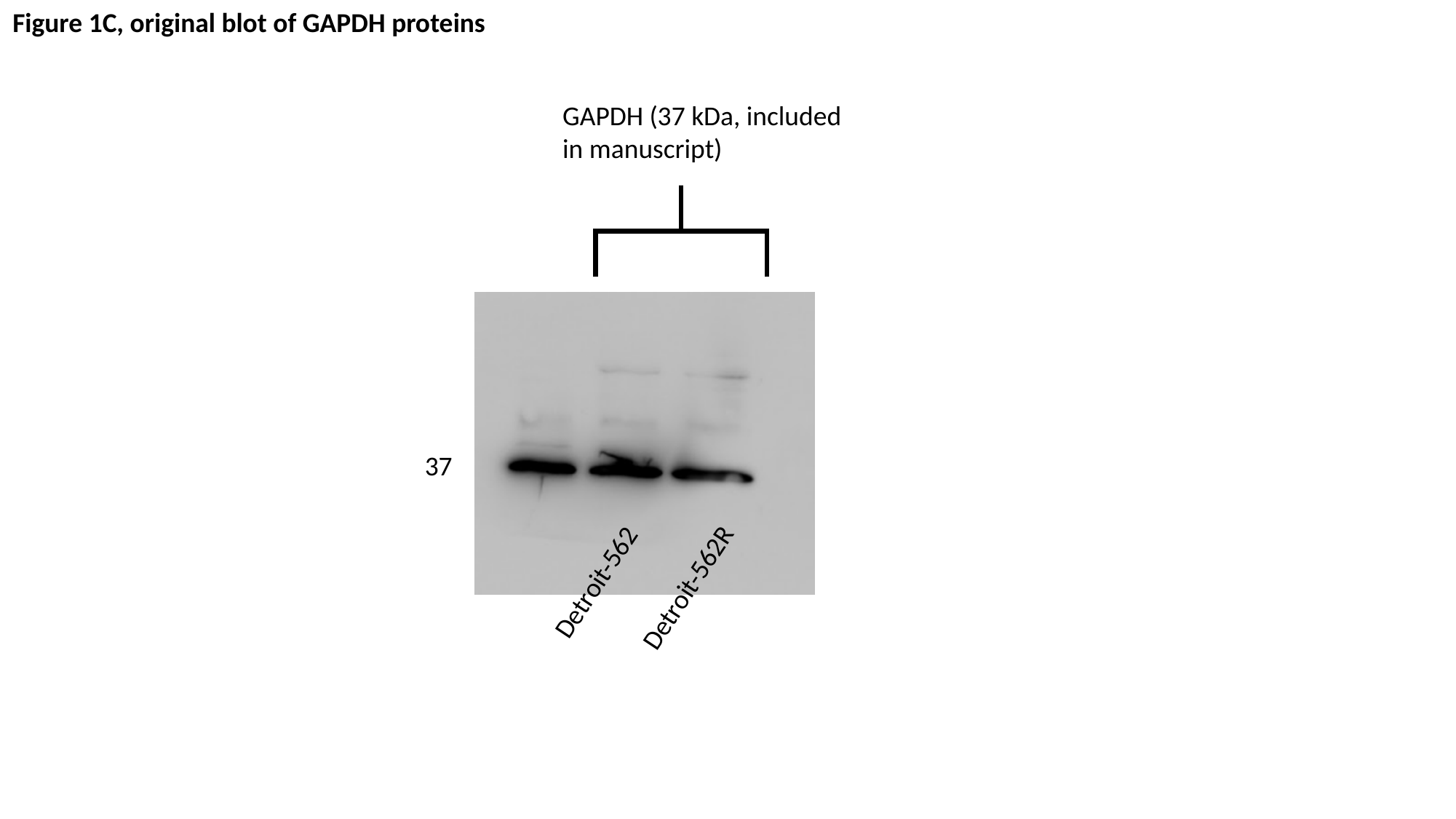

Figure 1C, original blot of GAPDH proteins
GAPDH (37 kDa, included in manuscript)
37
Detroit-562
Detroit-562R

### Slide 9
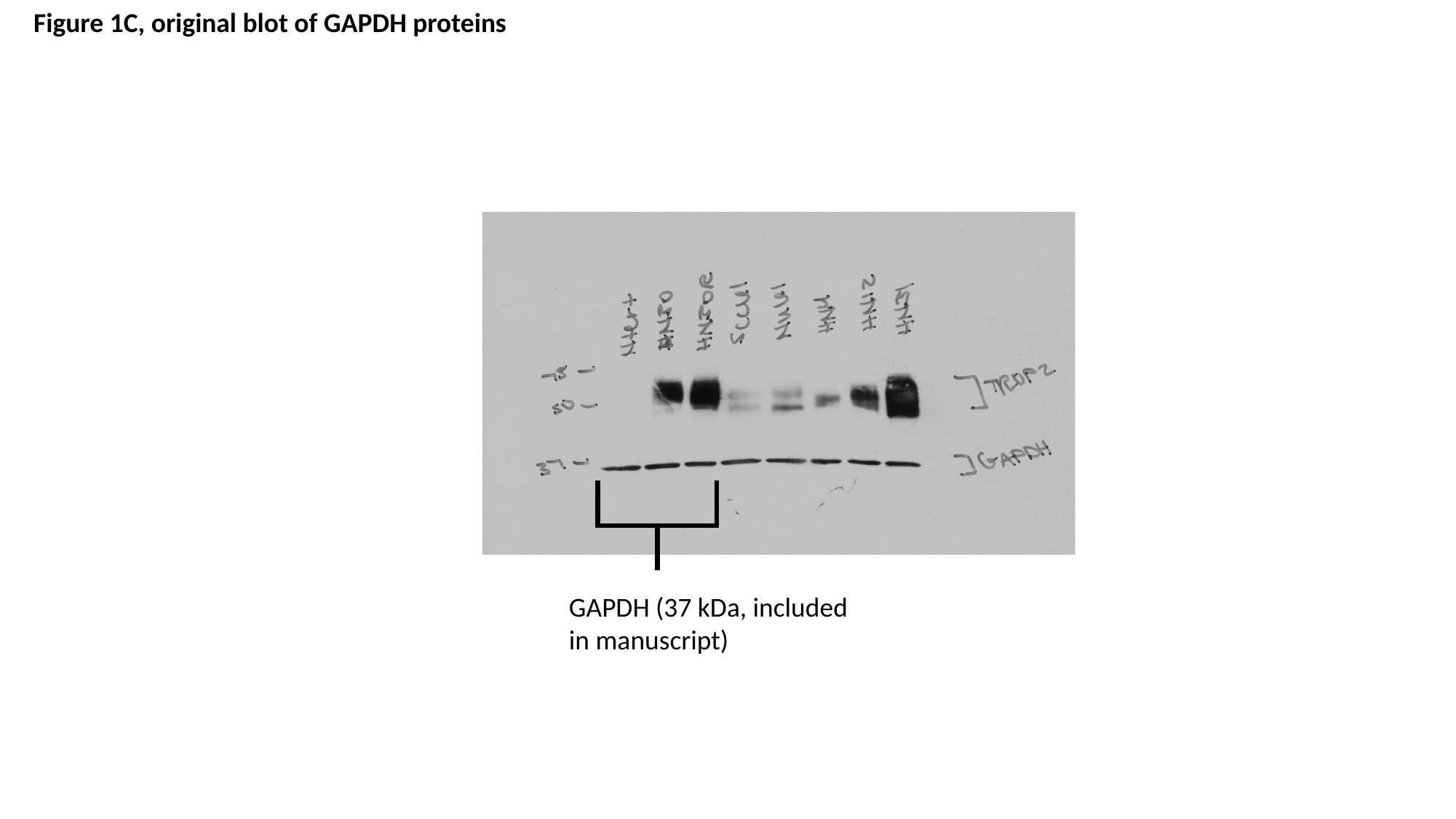

Figure 1C, original blot of GAPDH proteins
GAPDH (37 kDa, included in manuscript)

### Slide 10
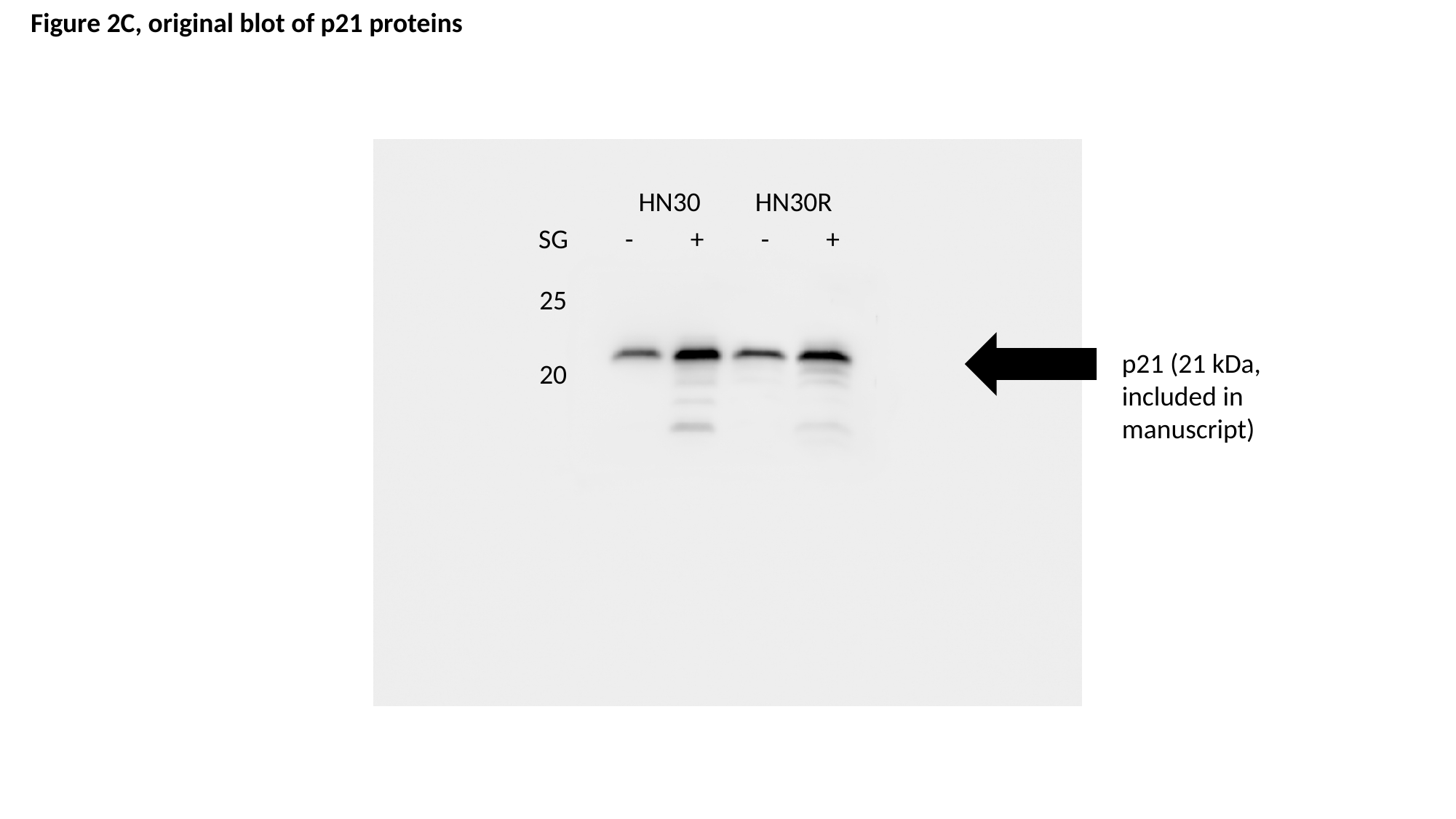

Figure 2C, original blot of p21 proteins
HN30
HN30R
| SG | - | + | - | + |
| --- | --- | --- | --- | --- |
25
p21 (21 kDa, included in manuscript)
20

### Slide 11
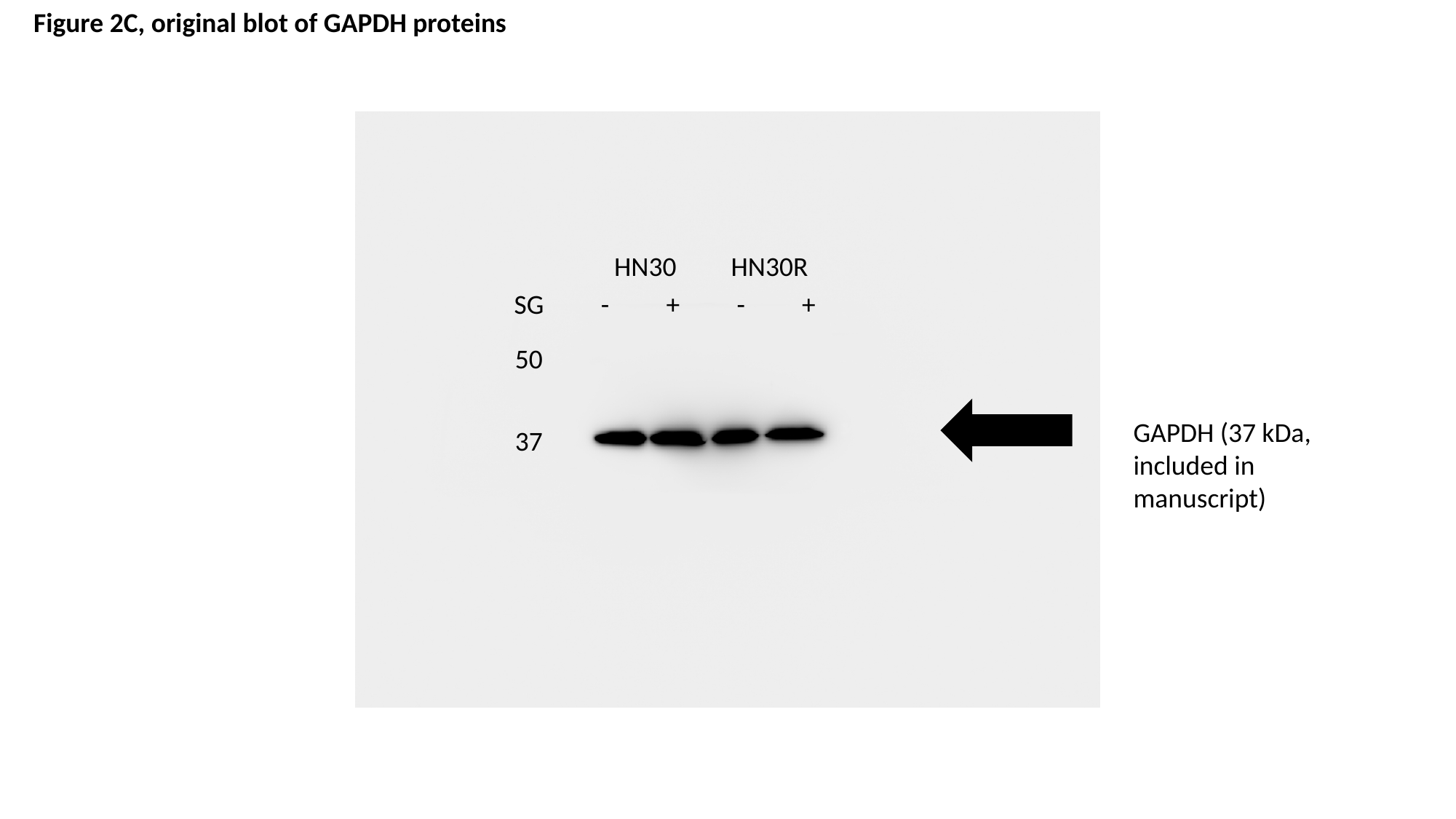

Figure 2C, original blot of GAPDH proteins
HN30
HN30R
| SG | - | + | - | + |
| --- | --- | --- | --- | --- |
50
GAPDH (37 kDa, included in manuscript)
37

### Slide 12
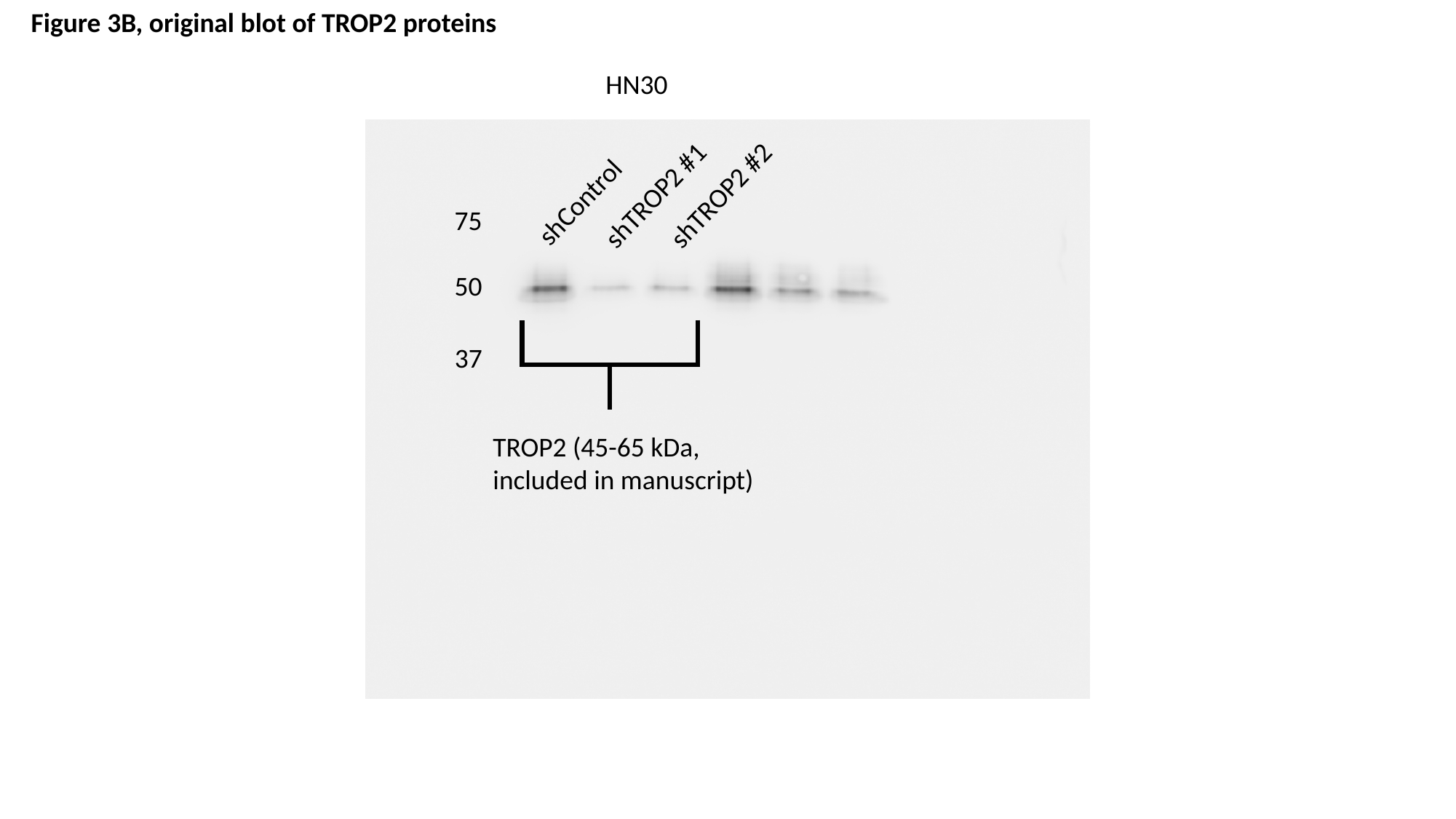

Figure 3B, original blot of TROP2 proteins
HN30
shTROP2 #1
shTROP2 #2
shControl
75
50
37
TROP2 (45-65 kDa, included in manuscript)

### Slide 13
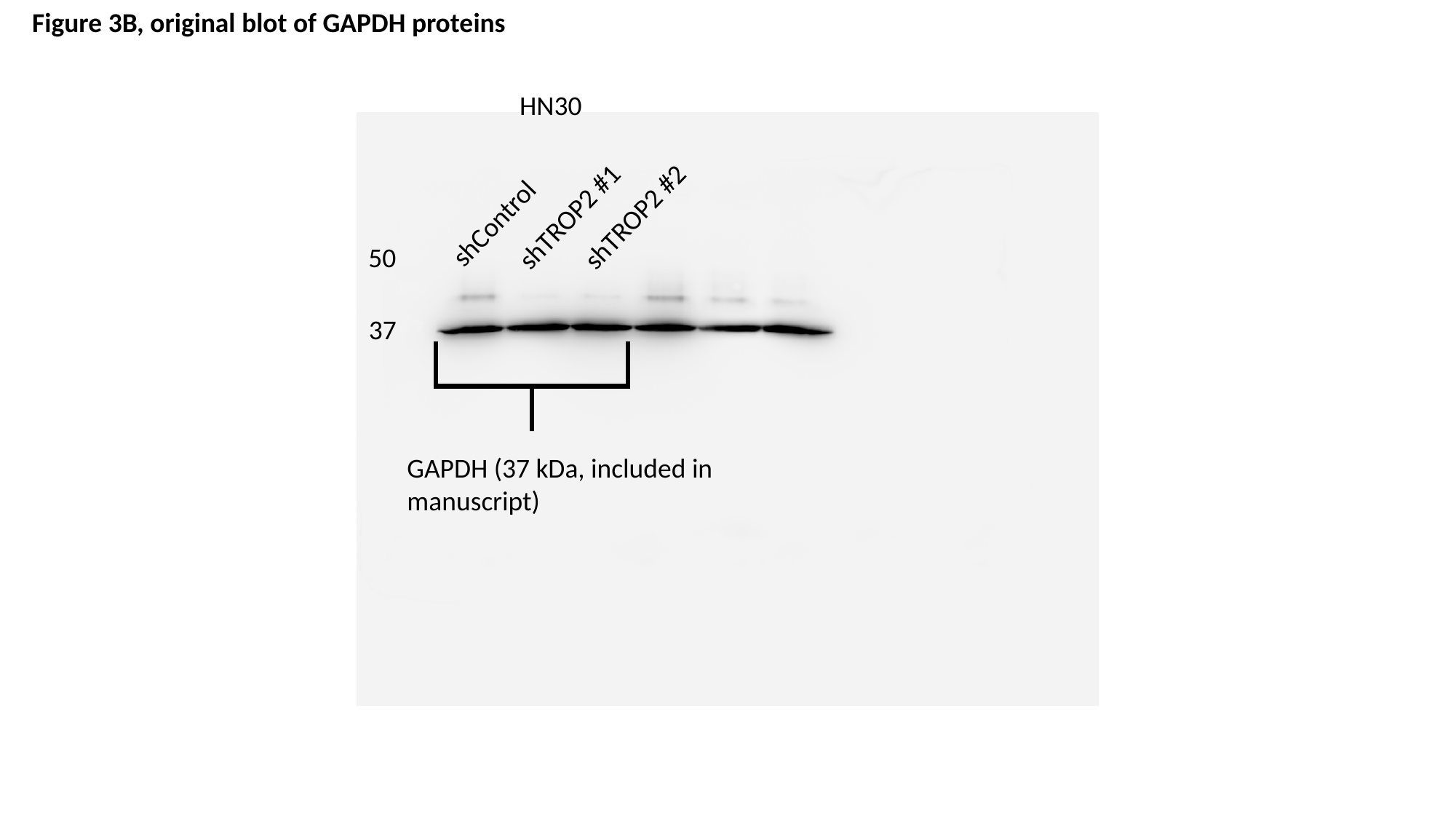

Figure 3B, original blot of GAPDH proteins
HN30
shTROP2 #1
shTROP2 #2
shControl
50
37
GAPDH (37 kDa, included in manuscript)

### Slide 14
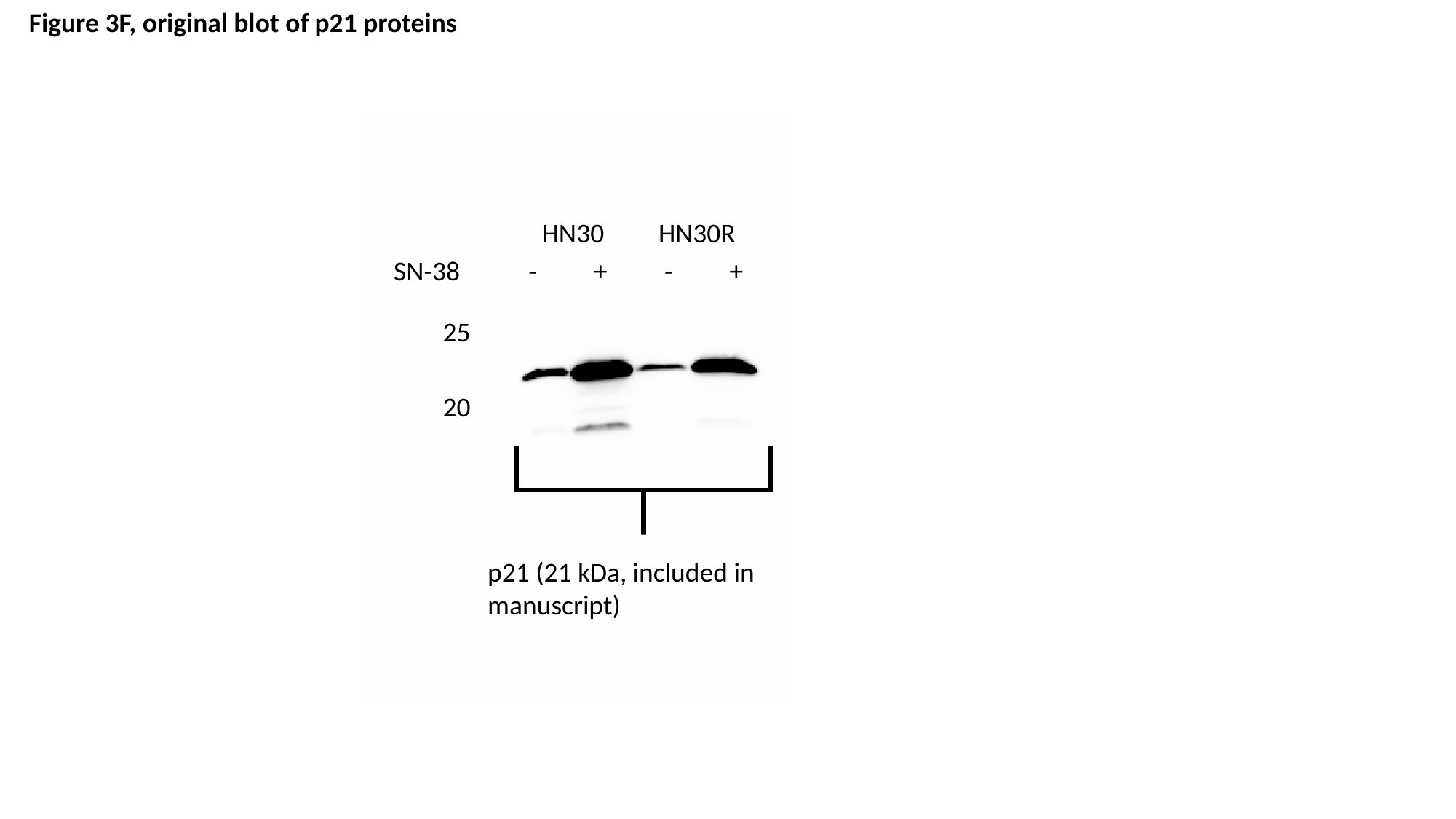

Figure 3F, original blot of p21 proteins
HN30
HN30R
| SN-38 | - | + | - | + |
| --- | --- | --- | --- | --- |
25
20
p21 (21 kDa, included in manuscript)

### Slide 15
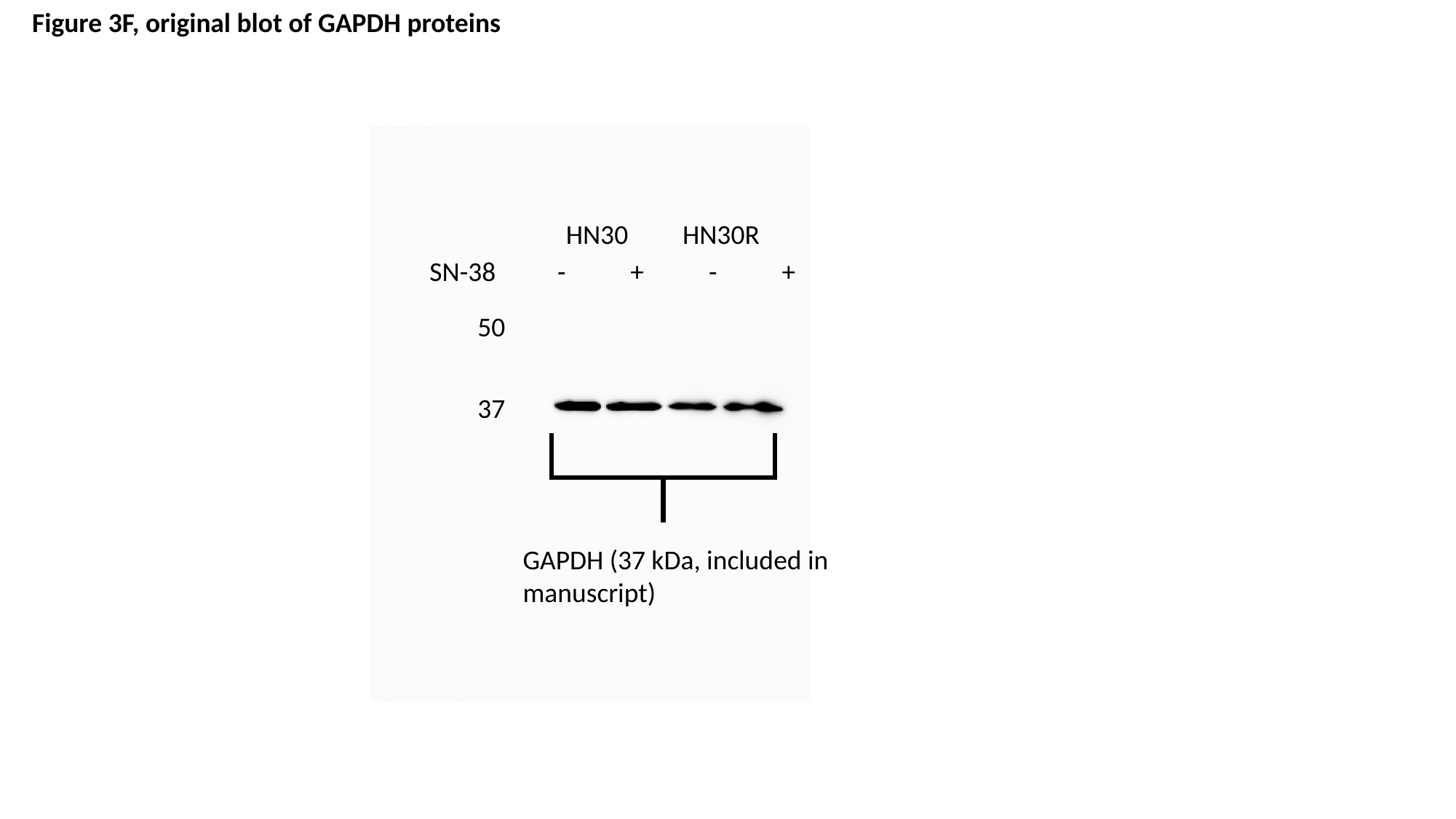

Figure 3F, original blot of GAPDH proteins
HN30
HN30R
| SN-38 | - | + | - | + |
| --- | --- | --- | --- | --- |
50
37
GAPDH (37 kDa, included in manuscript)

### Slide 16
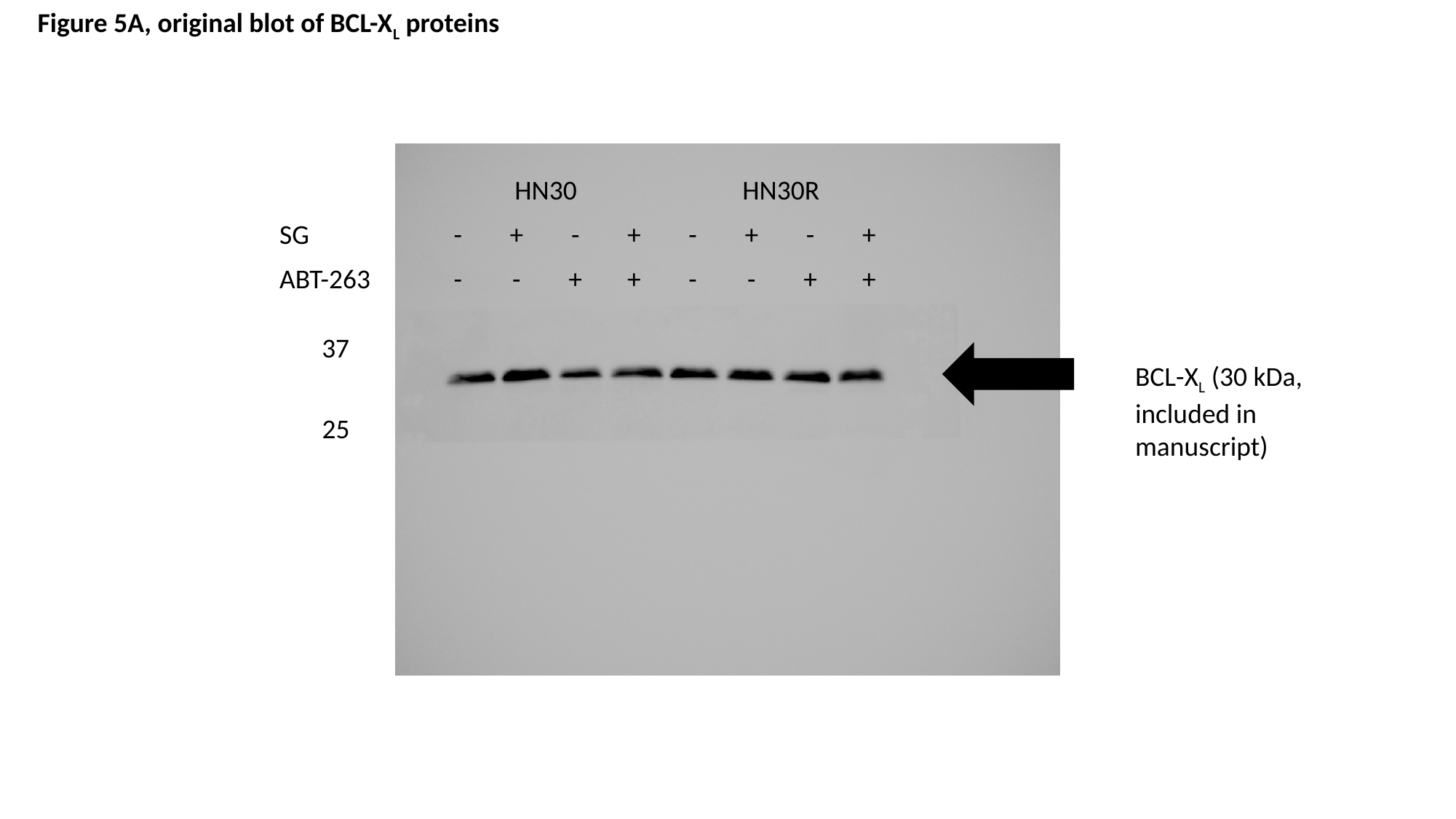

Figure 5A, original blot of BCL-X_L_ proteins
| | HN30 | | | | HN30R | | | |
| --- | --- | --- | --- | --- | --- | --- | --- | --- |
| SG | - | + | - | + | - | + | - | + |
| ABT-263 | - | - | + | + | - | - | + | + |
37
BCL-X_L_ (30 kDa, included in manuscript)
25

### Slide 17
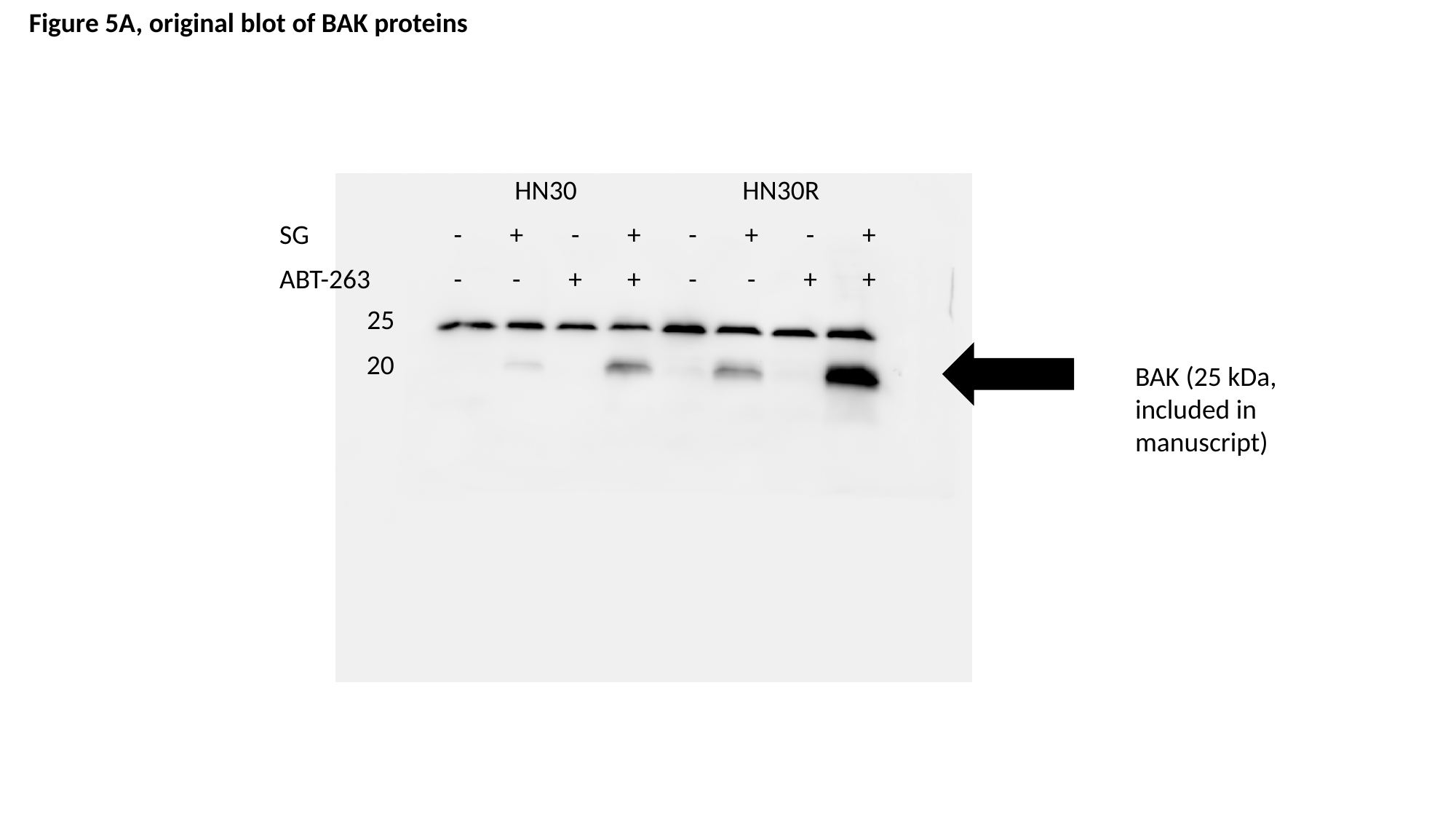

Figure 5A, original blot of BAK proteins
| | HN30 | | | | HN30R | | | |
| --- | --- | --- | --- | --- | --- | --- | --- | --- |
| SG | - | + | - | + | - | + | - | + |
| ABT-263 | - | - | + | + | - | - | + | + |
25
20
BAK (25 kDa, included in manuscript)

### Slide 18
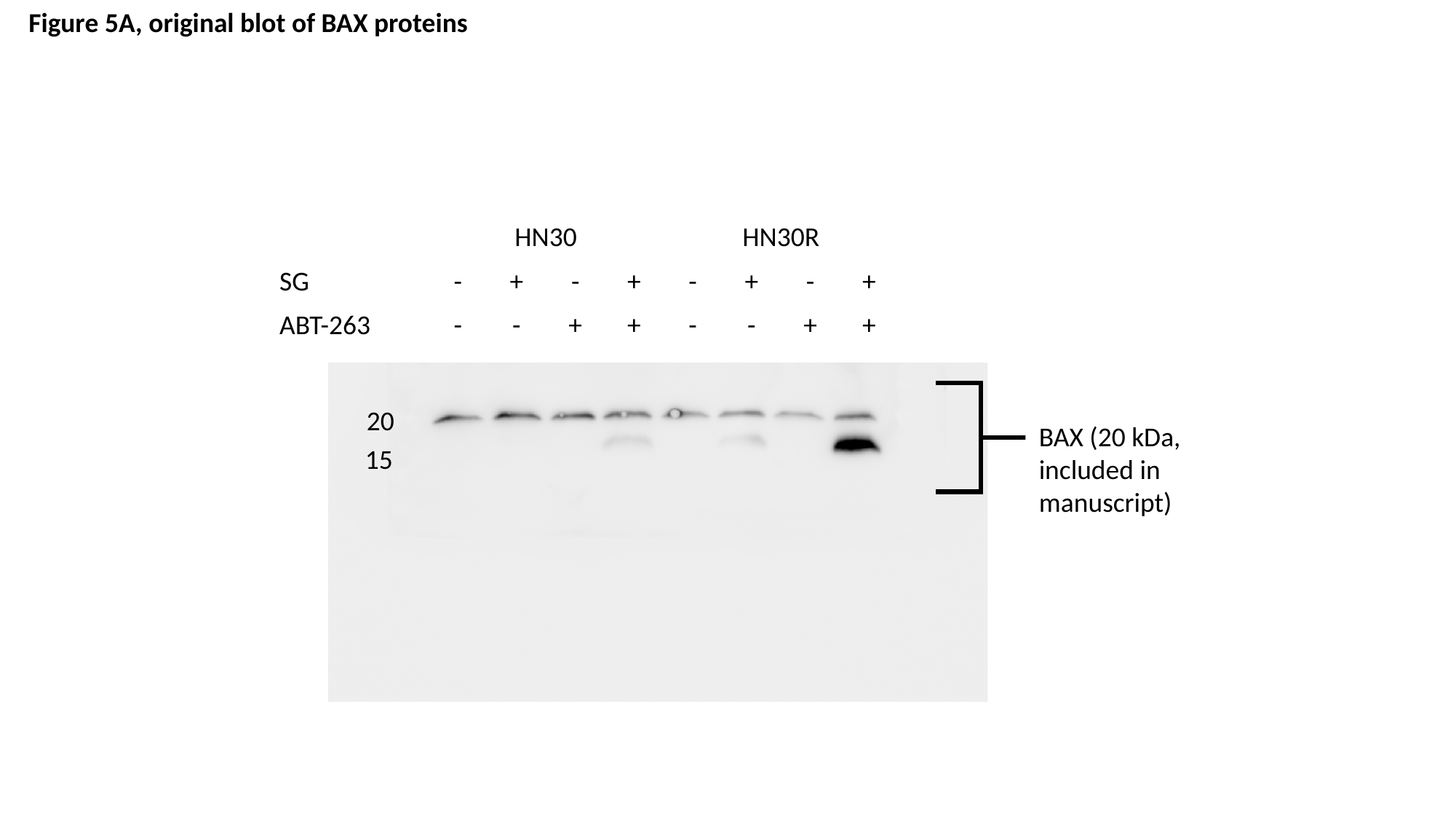

Figure 5A, original blot of BAX proteins
| | HN30 | | | | HN30R | | | |
| --- | --- | --- | --- | --- | --- | --- | --- | --- |
| SG | - | + | - | + | - | + | - | + |
| ABT-263 | - | - | + | + | - | - | + | + |
20
BAX (20 kDa, included in manuscript)
15

### Slide 19
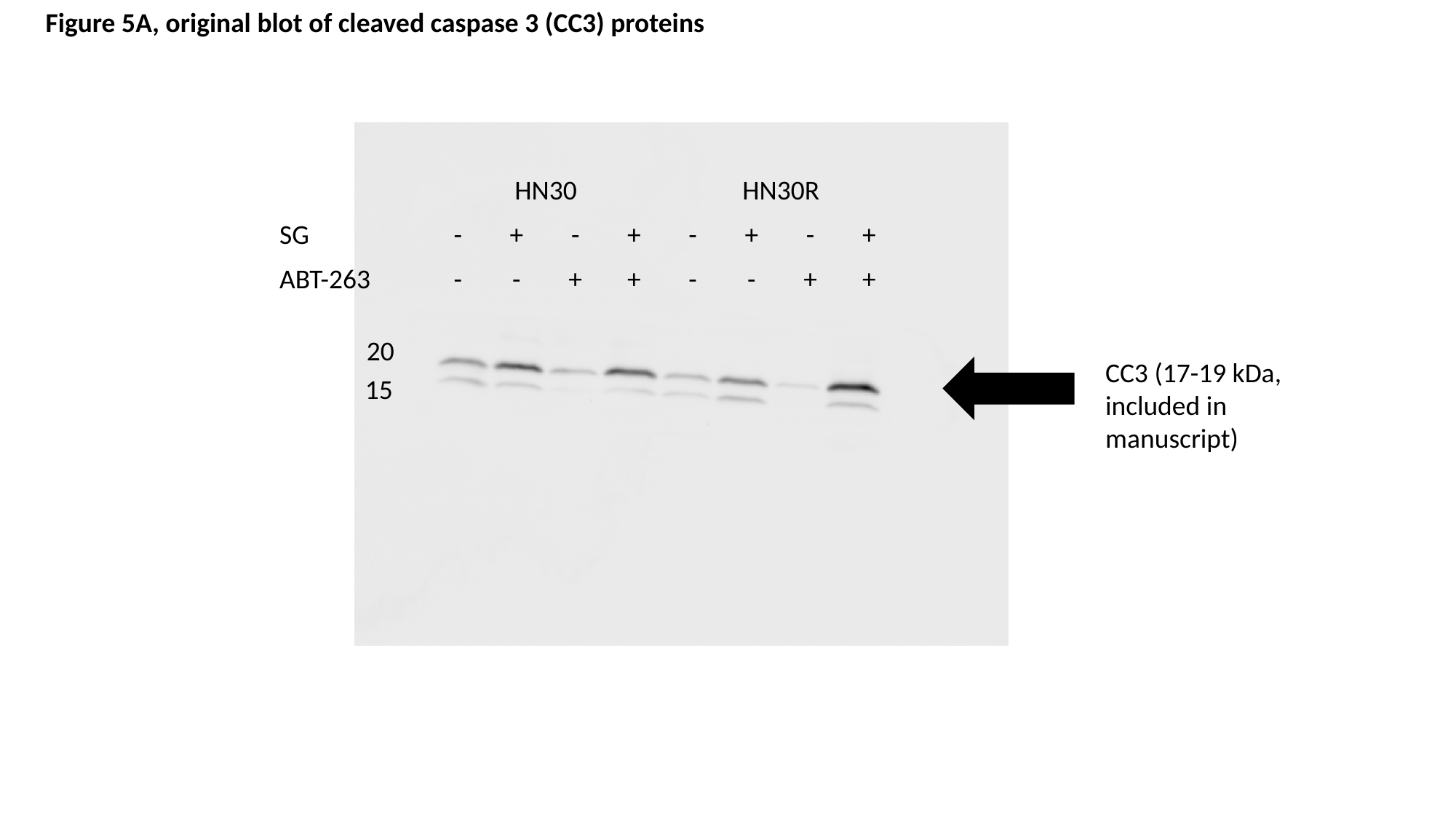

Figure 5A, original blot of cleaved caspase 3 (CC3) proteins
| | HN30 | | | | HN30R | | | |
| --- | --- | --- | --- | --- | --- | --- | --- | --- |
| SG | - | + | - | + | - | + | - | + |
| ABT-263 | - | - | + | + | - | - | + | + |
20
CC3 (17-19 kDa, included in manuscript)
15

### Slide 20
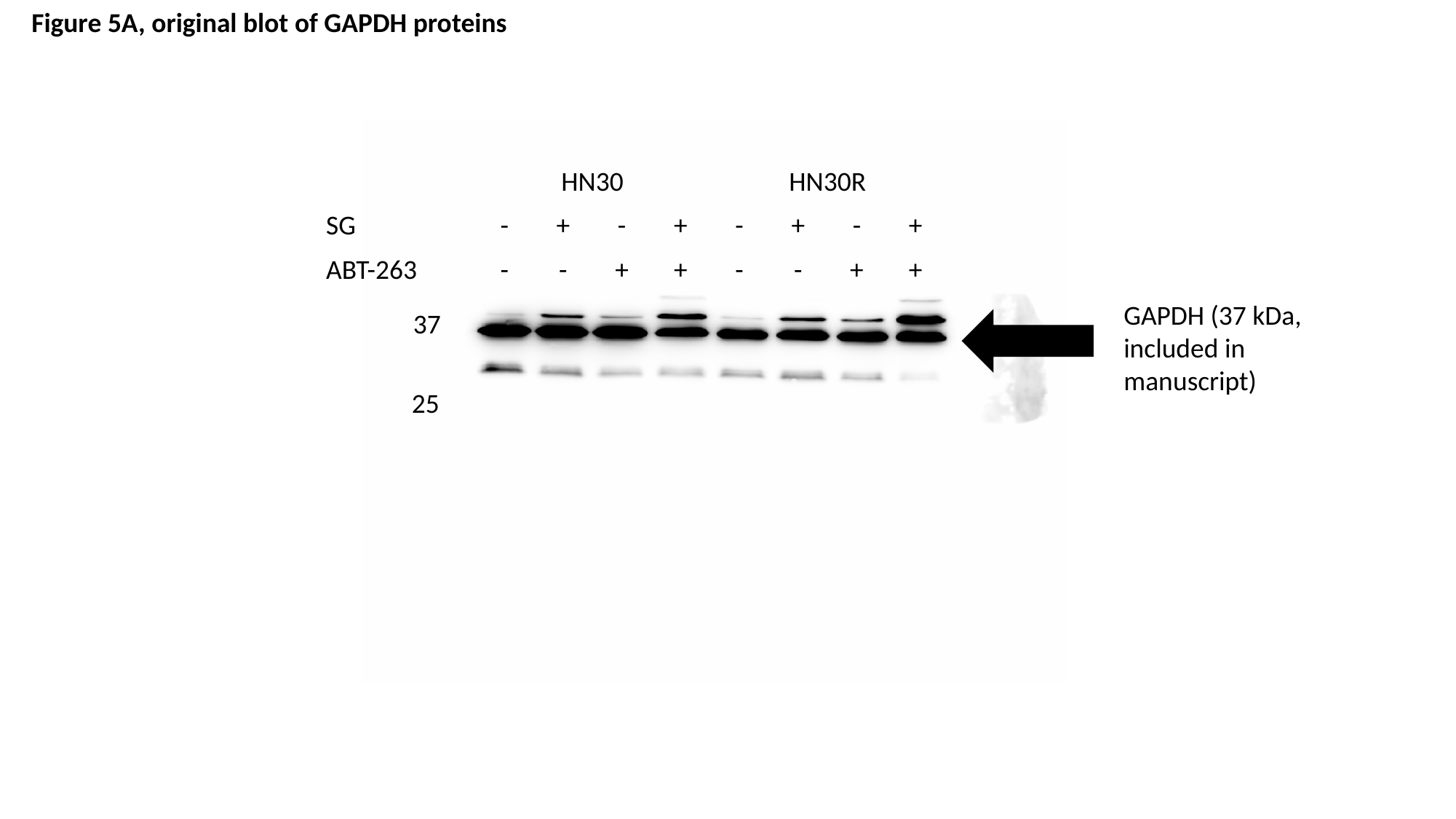

Figure 5A, original blot of GAPDH proteins
| | HN30 | | | | HN30R | | | |
| --- | --- | --- | --- | --- | --- | --- | --- | --- |
| SG | - | + | - | + | - | + | - | + |
| ABT-263 | - | - | + | + | - | - | + | + |
GAPDH (37 kDa, included in manuscript)
37
25

### Slide 21
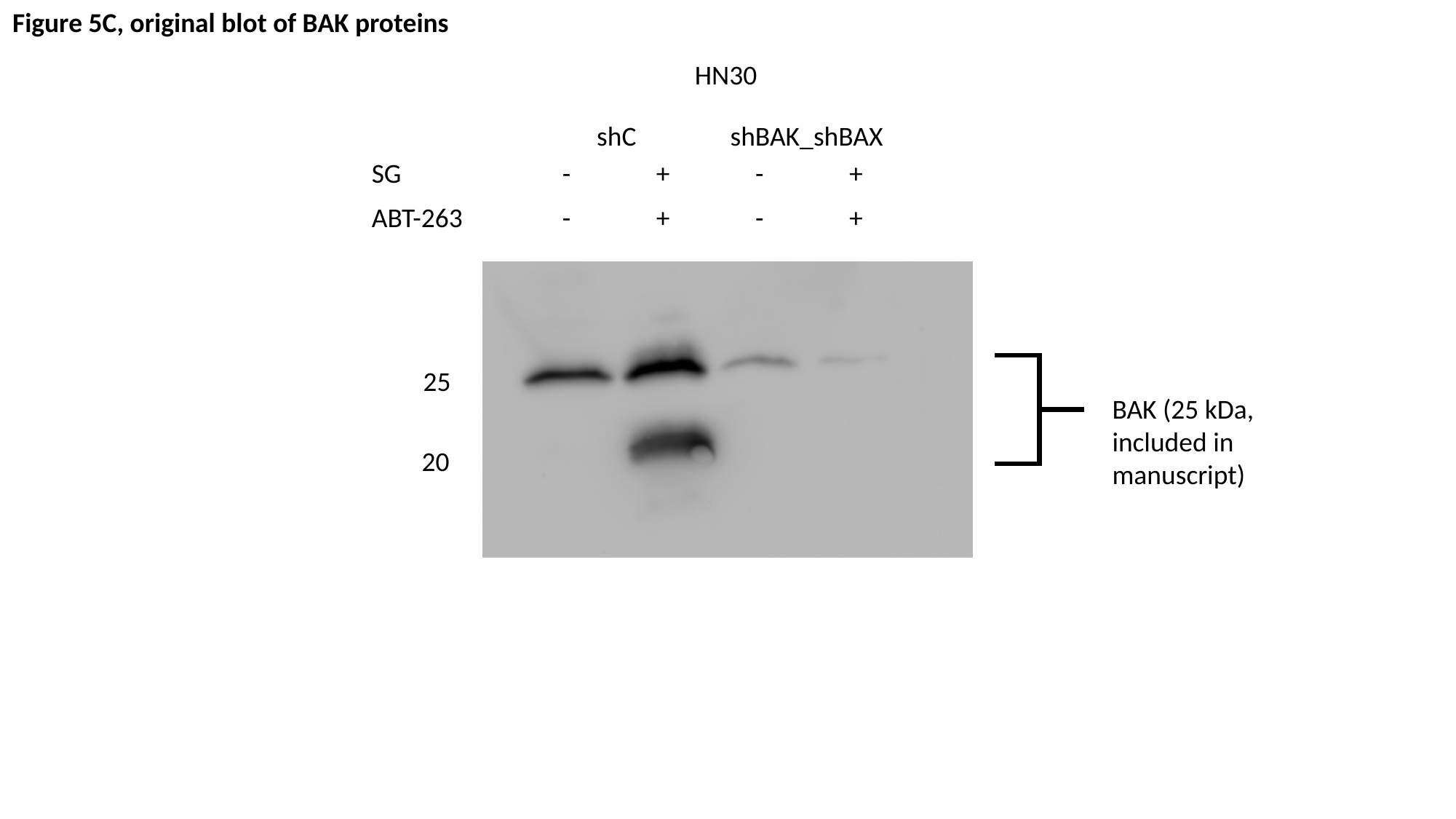

Figure 5C, original blot of BAK proteins
HN30
shC
shBAK_shBAX
| SG | - | + | - | + |
| --- | --- | --- | --- | --- |
| ABT-263 | - | + | - | + |
25
BAK (25 kDa, included in manuscript)
20

### Slide 22
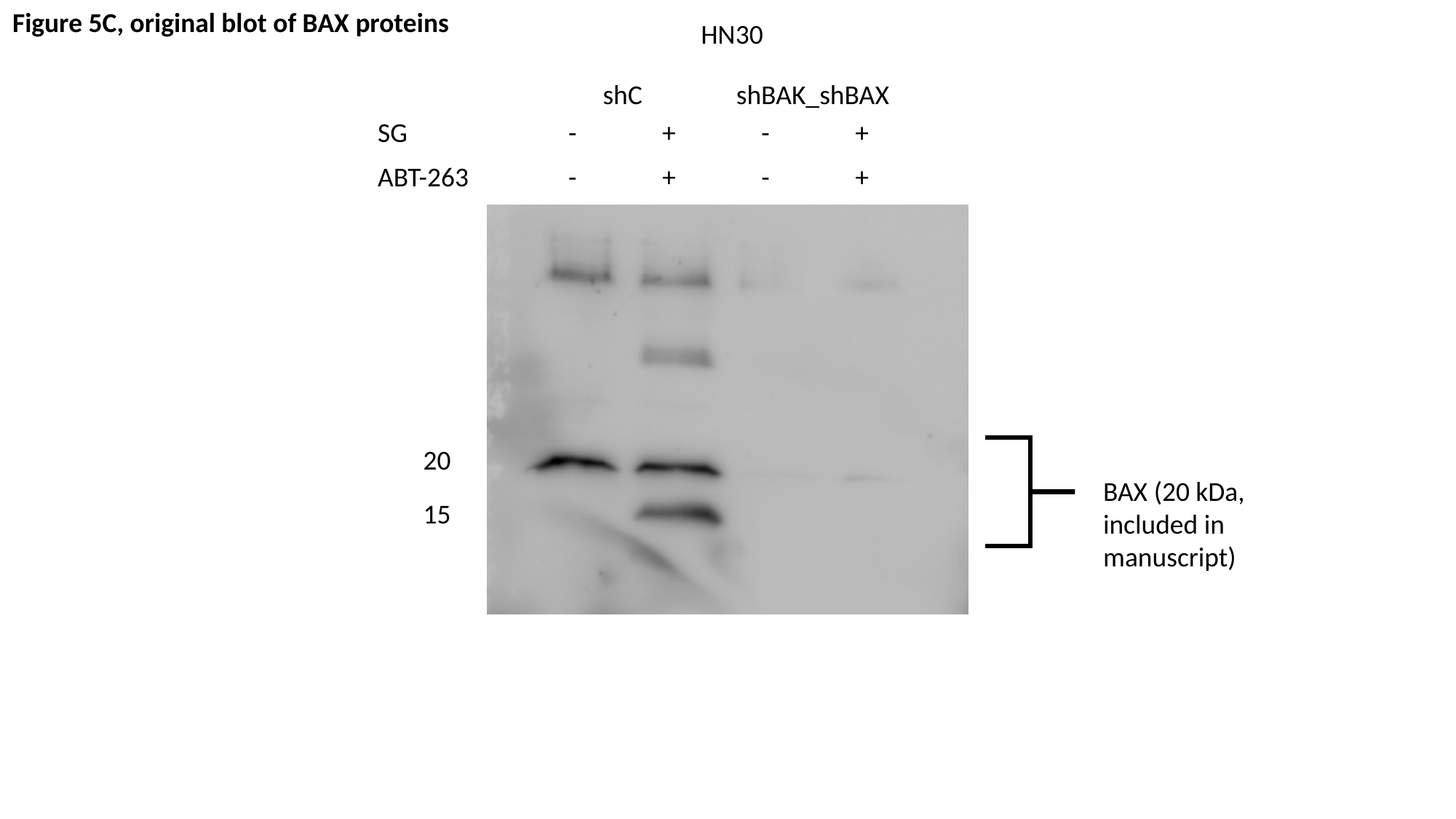

Figure 5C, original blot of BAX proteins
HN30
shC
shBAK_shBAX
| SG | - | + | - | + |
| --- | --- | --- | --- | --- |
| ABT-263 | - | + | - | + |
20
BAX (20 kDa, included in manuscript)
15

### Slide 23
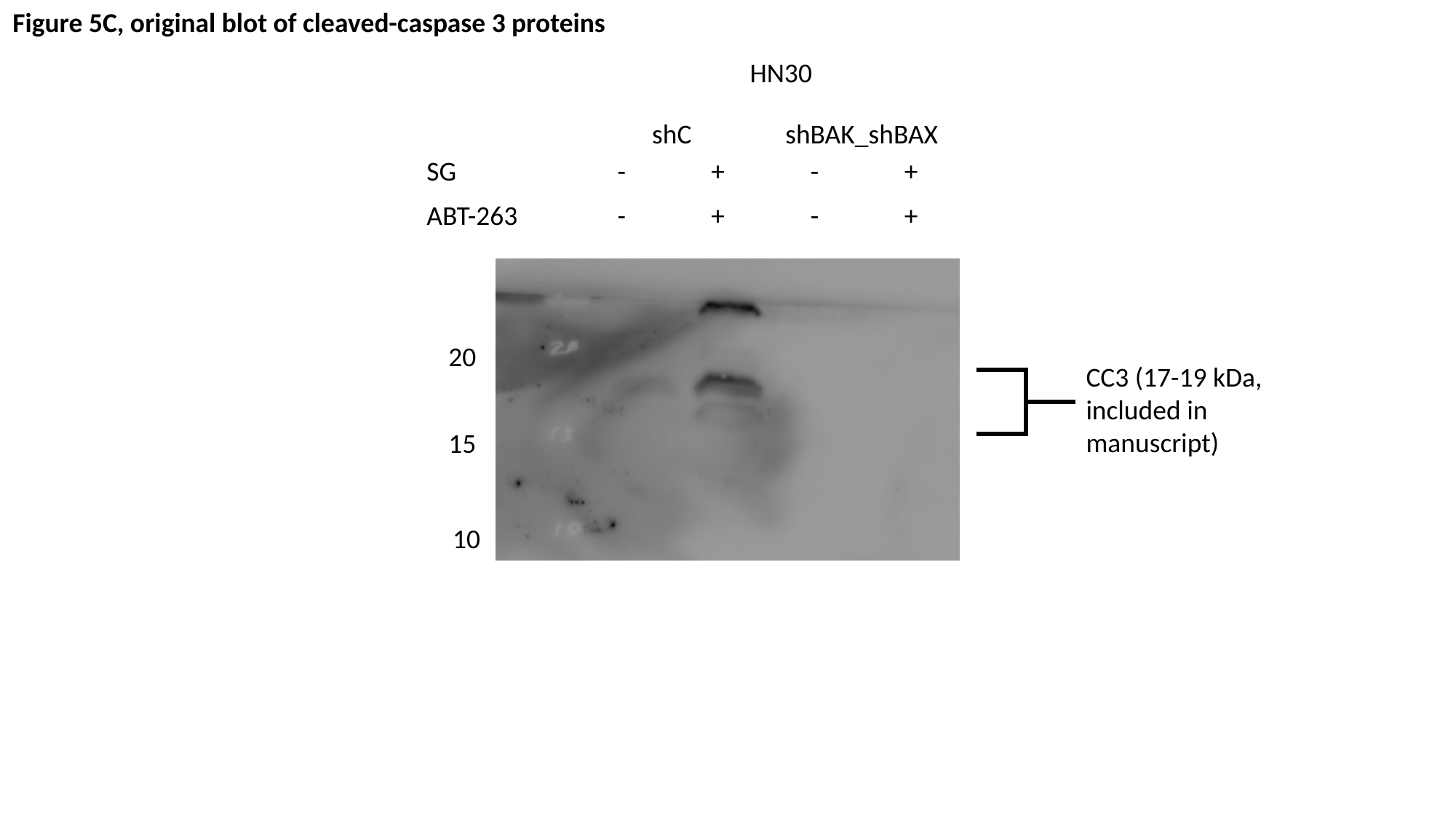

Figure 5C, original blot of cleaved-caspase 3 proteins
HN30
shC
shBAK_shBAX
| SG | - | + | - | + |
| --- | --- | --- | --- | --- |
| ABT-263 | - | + | - | + |
20
CC3 (17-19 kDa, included in manuscript)
15
10

### Slide 24
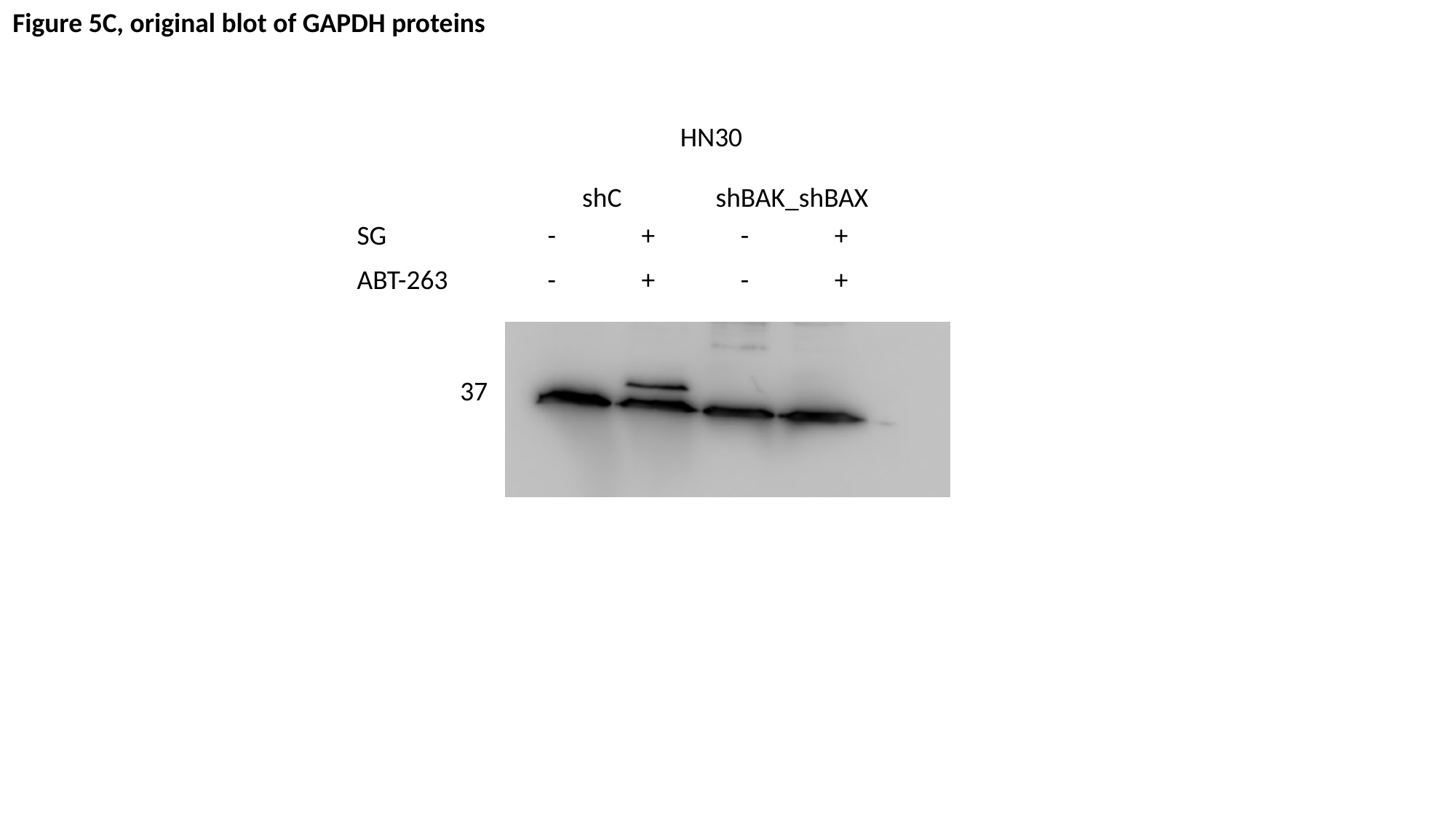

Figure 5C, original blot of GAPDH proteins
HN30
shC
shBAK_shBAX
| SG | - | + | - | + |
| --- | --- | --- | --- | --- |
| ABT-263 | - | + | - | + |
37

### Slide 25
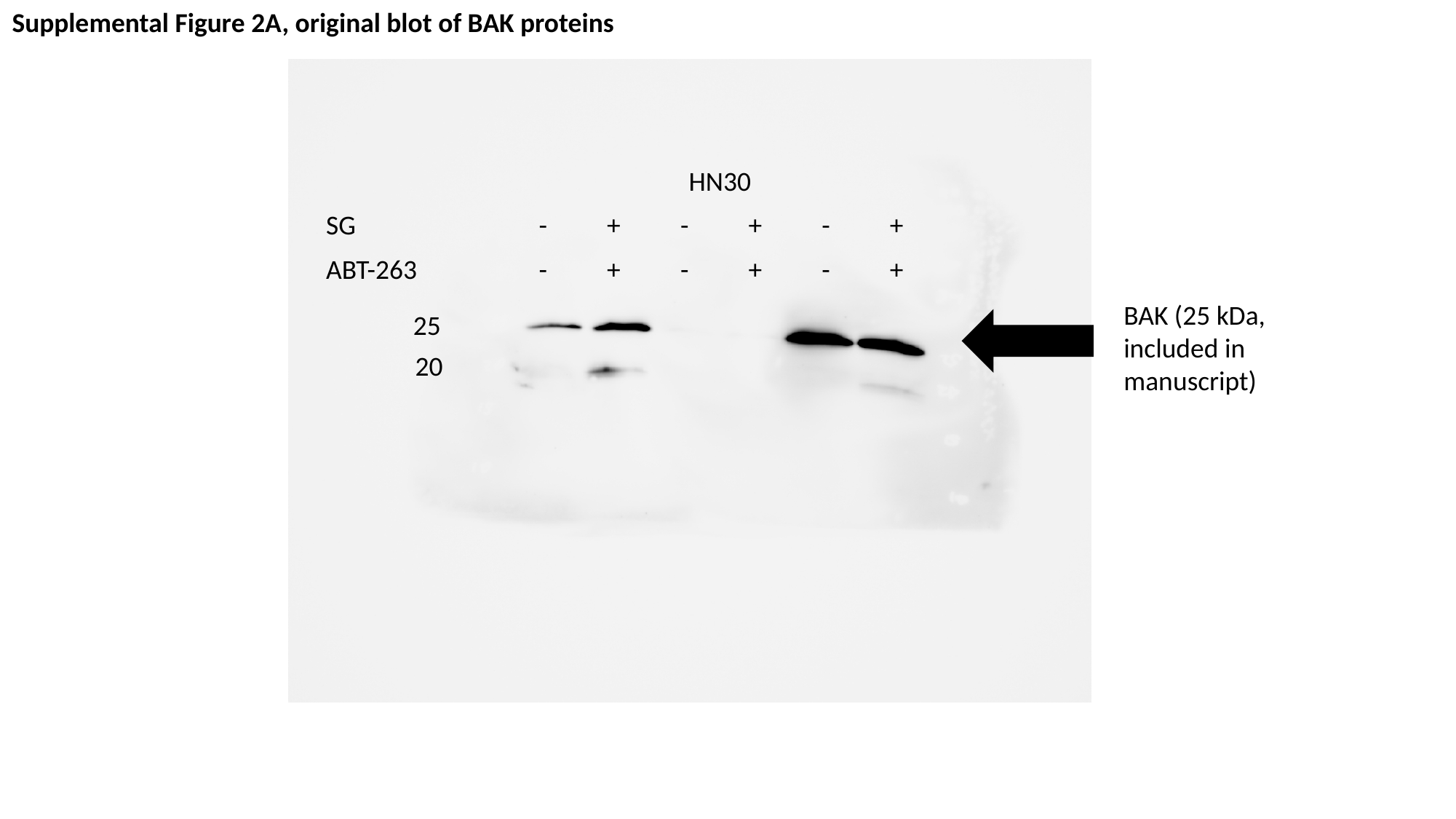

Supplemental Figure 2A, original blot of BAK proteins
| | HN30 | | | | | |
| --- | --- | --- | --- | --- | --- | --- |
| SG | - | + | - | + | - | + |
| ABT-263 | - | + | - | + | - | + |
BAK (25 kDa, included in manuscript)
25
20

### Slide 26
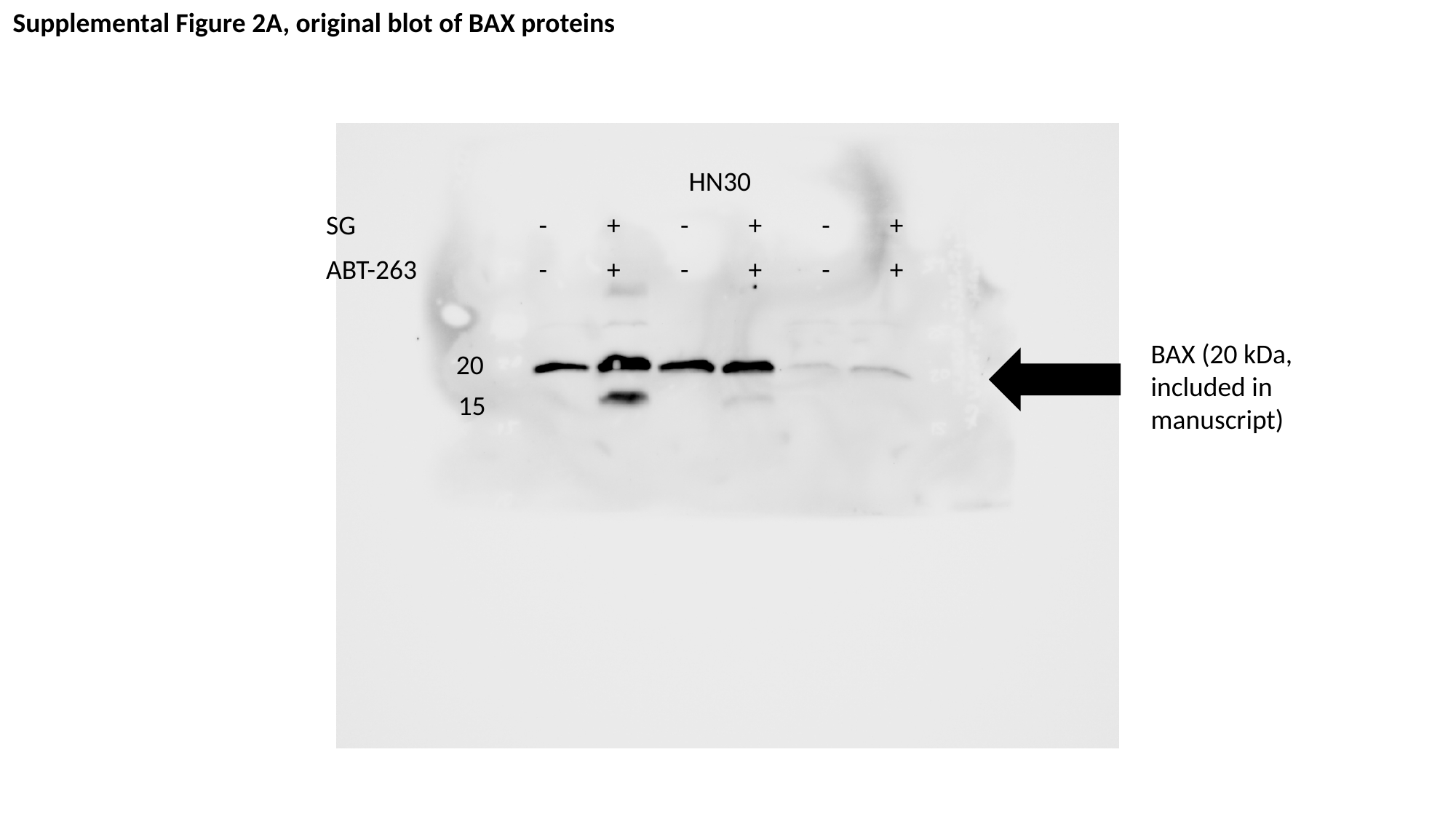

Supplemental Figure 2A, original blot of BAX proteins
| | HN30 | | | | | |
| --- | --- | --- | --- | --- | --- | --- |
| SG | - | + | - | + | - | + |
| ABT-263 | - | + | - | + | - | + |
BAX (20 kDa, included in manuscript)
20
15

### Slide 27
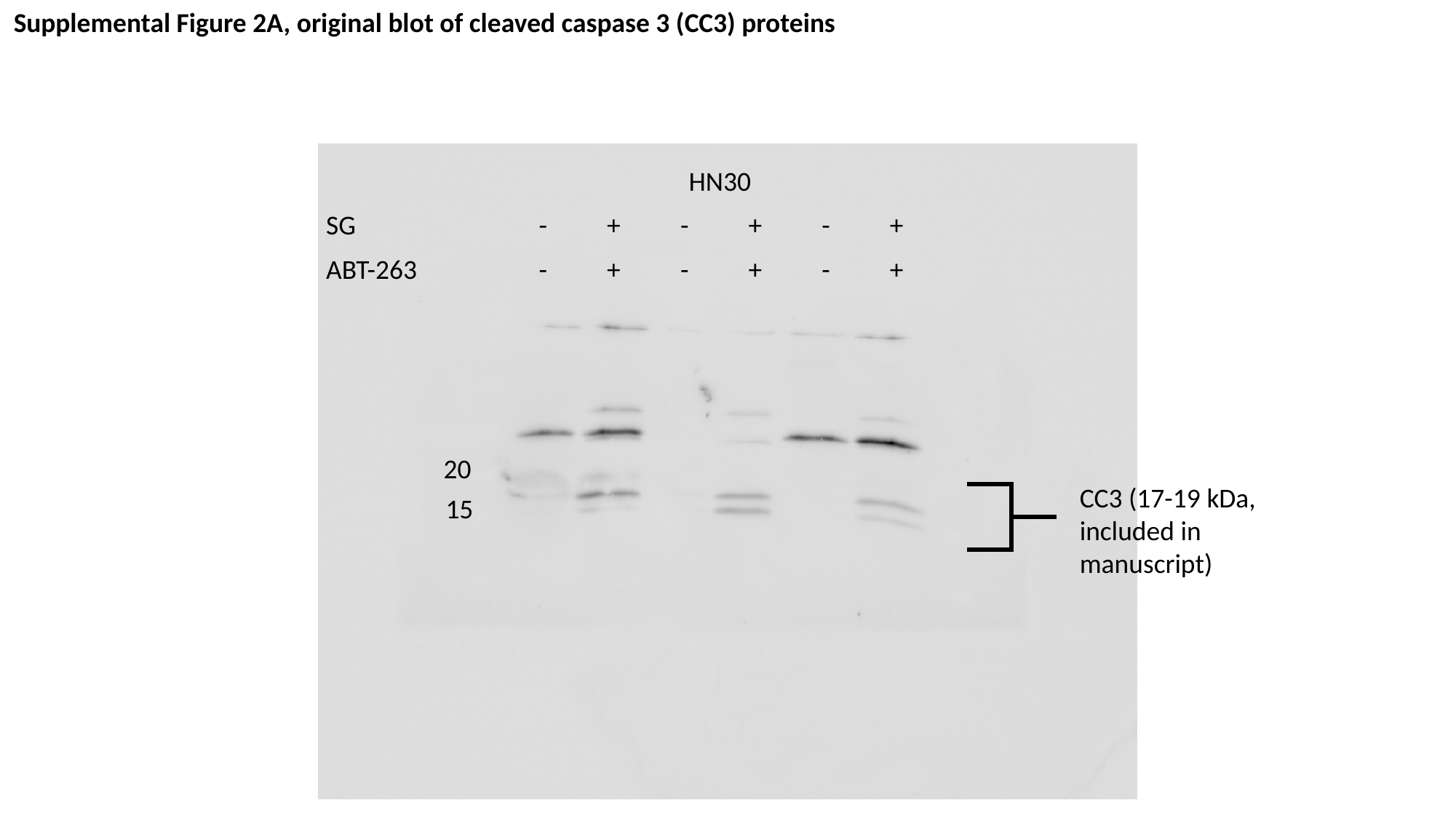

Supplemental Figure 2A, original blot of cleaved caspase 3 (CC3) proteins
| | HN30 | | | | | |
| --- | --- | --- | --- | --- | --- | --- |
| SG | - | + | - | + | - | + |
| ABT-263 | - | + | - | + | - | + |
20
CC3 (17-19 kDa, included in manuscript)
15

### Slide 28
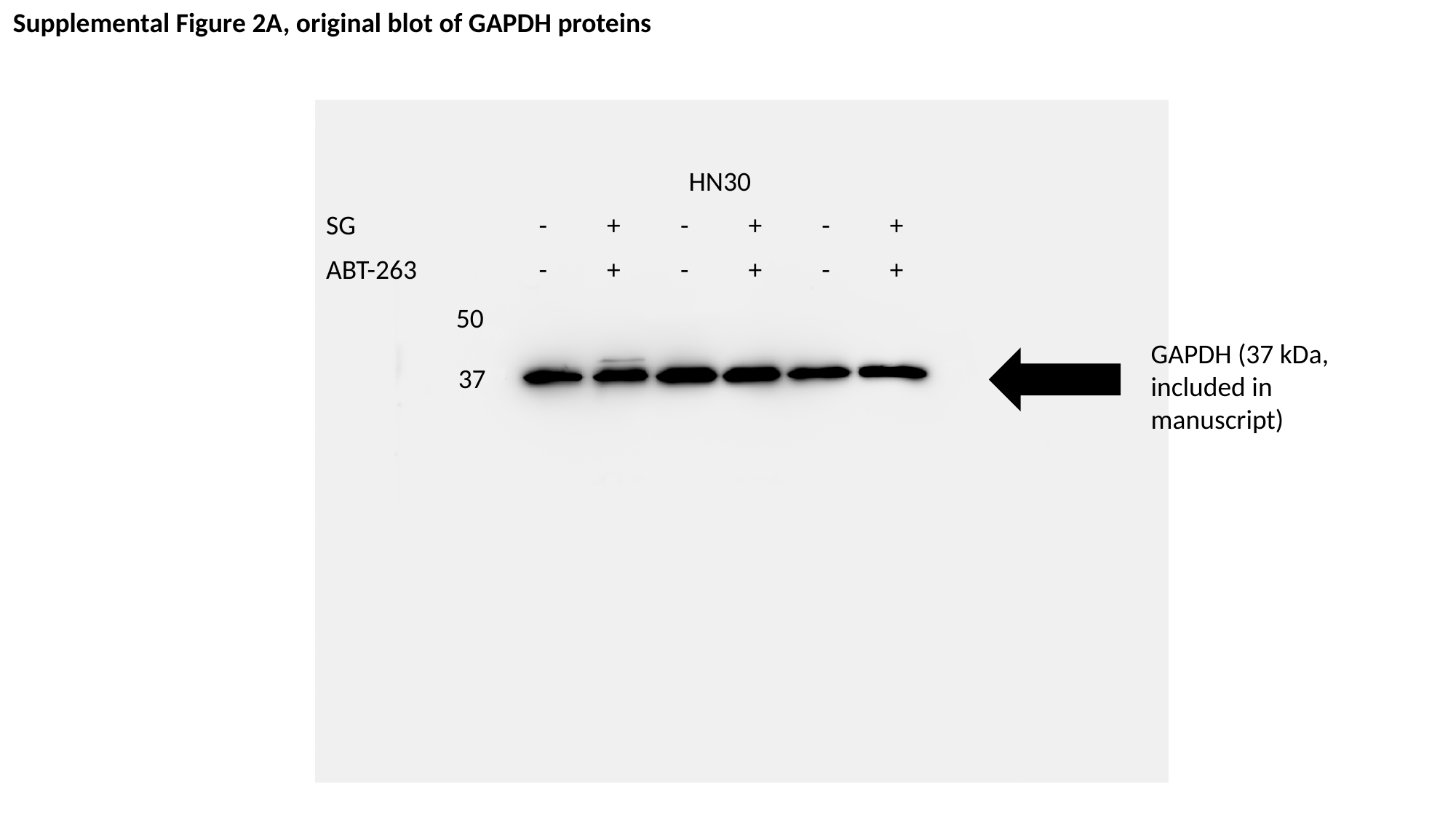

Supplemental Figure 2A, original blot of GAPDH proteins
| | HN30 | | | | | |
| --- | --- | --- | --- | --- | --- | --- |
| SG | - | + | - | + | - | + |
| ABT-263 | - | + | - | + | - | + |
50
GAPDH (37 kDa, included in manuscript)
37
